# Circulating multimeric immune complexes drive immunopathology in COVID-19

**DOI:** 10.1101/2021.06.25.449893

**Authors:** Jakob Ankerhold, Sebastian Giese, Philipp Kolb, Andrea Maul-Pavicic, Reinhard E. Voll, Nathalie Göppert, Kevin Ciminski, Clemens Kreutz, Achim Lother, Ulrich Salzer, Wolfgang Bildl, Tim Welsink, Nils G. Morgenthaler, Andrea Busse Grawitz, Daniela Huzly, Martin Schwemmle, Hartmut Hengel, Valeria Falcone

## Abstract

A dysregulated immune response with high levels of SARS-CoV-2 specific IgG antibodies characterizes patients with severe or critical COVID-19. Although a robust IgG response is traditionally considered to be protective, excessive triggering of activating Fc-gamma-receptors (FcγRs) could be detrimental and cause immunopathology. Here, we document that patients who develop soluble circulating IgG immune complexes (sICs) during infection are subject to enhanced immunopathology driven by FcγR activation. Utilizing cell-based reporter systems we provide evidence that sICs are predominantly formed prior to a specific humoral response against SARS-CoV-2. sIC formation, together with increased afucosylation of SARS-CoV-2 specific IgG eventually leads to an enhanced CD16 (FcγRIII) activation of immune cells reaching activation levels comparable active systemic lupus erythematosus (SLE) disease. Our data suggest a vicious cycle of escalating immunopathology driven by an early formation of sICs in predisposed patients. These findings reconcile the seemingly paradoxical findings of high antiviral IgG responses and systemic immune dysregulation in severe COVID-19.

**Clinical implications:** The identification of sICs as drivers of an escalating immunopathology in predisposed patients opens new avenues regarding intervention strategies to alleviate critical COVID-19 progression.

**Graphical abstract:** 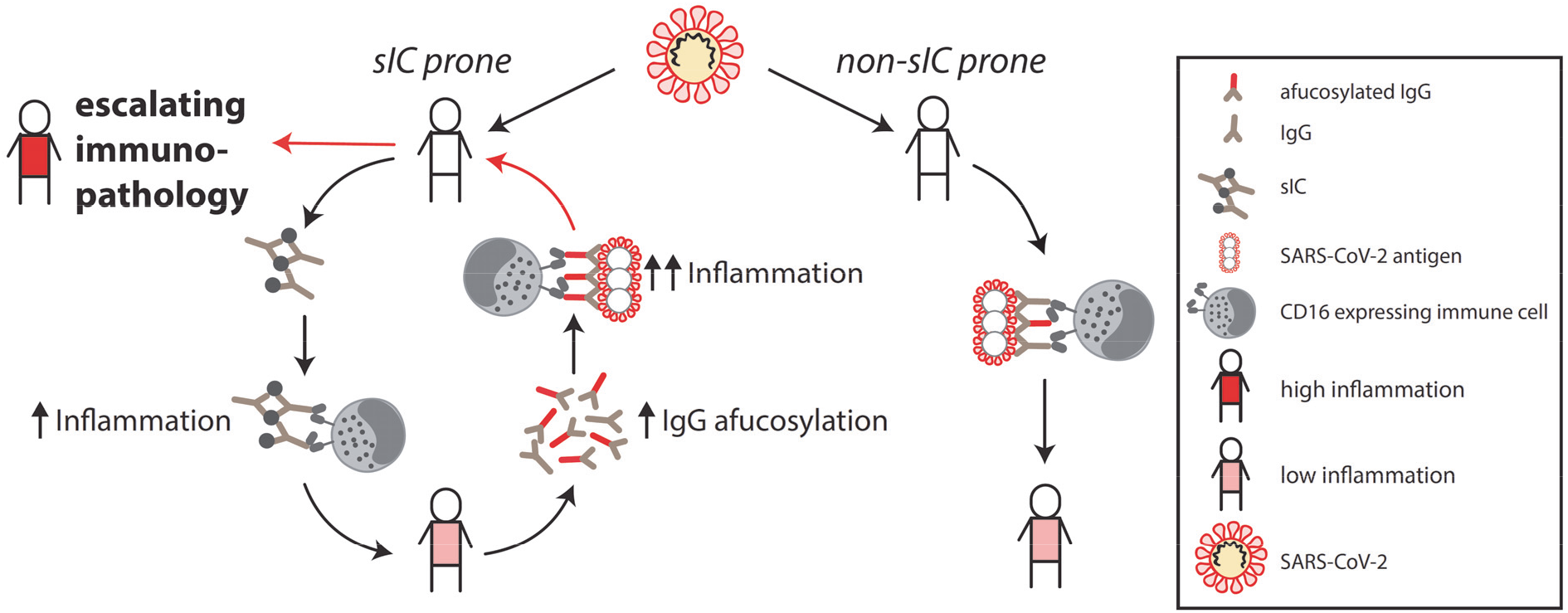

**A vicious cycle of immunopathology in COVID-19 patients is driven by soluble multimeric immune complexes (sICs)**. SARS-CoV-2 infection triggers sIC formation in prone individuals. Activation of FcγRIII/CD16 expressing immune cells by sICs precedes a humoral response to SARS-CoV2 infection. sICs and infection add to IgG afucosylation, further enhancing FcγRIII/CD16 activation by opsonized targets. High inflammation induces further sIC mediated immune cell activation ultimately leading to an escalating immunopathology.

## Introduction

Since the emergence of SARS-CoV-2 in late December 2019 (1), more than 199 million laboratory confirmed infections (as of August 3rd, 2021) have been reported, with cases continuously rising (2). Accordingly, rapid insights into the disease manifestations and pathogenesis have been globally obtained. A hallmark of the coronavirus disease 2019 (COVID-19) is a respiratory infection which can progress to an acute respiratory distress syndrome (ARDS). Next to asymptomatic infections, COVID-19 symptoms differ widely according to the disease process and may comprise fever, coughing, pneumonia, dyspnea, hypoxia and lymphopenia (3). While fever and coughing are common symptoms, pneumonia, hypoxia, dyspnea, certain organ manifestations and lymphopenia indicate critical or fatal infections (3-6). Pronounced dyspnea can eventually progress to ARDS, a severe complication frequently observed in critically ill patients (7, 8). Although overall disease severity and in particular breathing difficulties are related to viral load (9), age (4, 10-13) and underlying medical conditions (4, 11, 12), the delayed kinetics of respiratory failure strongly suggest an essential role of the host immune response (3, 11). Typically, aggravation occurs between 9-11 days after symptom onset (12) and correlates with high levels of SARS-CoV-2 specific IgG antibodies and systemic effects of pro-inflammatory cytokines such as IL-6 and TNFα (3, 14-16). This cytokine release, primarily the result of macrophage and T helper (T_H_) cell activation (17), includes pattern recognition receptor (PRR) signaling in the context of innate immunity but can also occur by Fcγ receptor (FcγR) activation (18). Triggered by immune complexes (antibody-antigen complex), the cytokine release following FcγR activation represents a potent defense mechanism against invading pathogens. A prototypical activating FcγR in this regard is FcγRIII (CD16) expressed by NK cells (19, 20), monocyte-derived macrophages (CD16A) (21) or neutrophils (CD16B, 98% sequence identical ectodomains). Specifically, CD16 is able to sense circulating soluble immune complexes (sICs) as they are formed in certain autoimmune diseases such as systemic lupus erythematosus (SLE) (22-25) and viral infections (26). Overstimulation of activating FcγRs in these cases is associated with disease severity (26-28) and thus an FcγR-driven overshooting inflammatory response (18) might be an explanation for the pronounced immunopathology observed during severe courses of COVID-19 (29). Consistently, hyper-inflammation in SARS-CoV-1 and MERS infected patients has been previously proposed as a possible pathogenic factor (30) and could be demonstrated in mice and macaques infected with SARS-CoV-1 (31, 32). Furthermore, N297-dependent glycan-modifications such as afucosylation within the constant region of IgG antibodies are known to enhance FcγR binding, in turn promoting inflammation. It has been shown that enhanced FcγRIII activation by low-fucosylated anti-SARS-CoV-2-S IgG leads to excessive alveolar macrophage activation, driving severe COVID-19 disease progression (33). Further, it has been proposed that uncleared antigen-antibody immune complexes (ICs) might be involved in the pathogenesis of severe disease leading to systemic complement activation and tissue damage, neutrophil activation, cytokine storm, systemic vasculitis, microvascular thrombosis and organ failure (34-39). However, comprehensive evidence that circulating sICs impact disease progression is still missing. Here, we aimed to further delineate the contribution of IgG-mediated effector functions regarding COVID-19 severity in patient cohorts with varying disease progressions. This revealed a marked correlation between CD16 activation by patient IgG and severity of disease. Additionally, we identified circulating CD16-reactive sICs to be abundantly present in the serum of patients with critical and severe disease, but not in the serum of patients with a mild disease. sIC levels were comparable to those found in SLE patients with active disease. As sIC formation preceded a SARS-CoV-2 specific humoral response in most cases, we conclude that a so far undisclosed predisposing condition divides patients into sIC-prone and non-sIC-prone individuals with patients developing sICs in response to an infectious trigger also developing enhanced disease. Our data suggest a vicious cycle leading to an escalating immunopathology driven by the early formation of sICs. Our findings enable new avenues of intervention against COVID-19 and highly warrant further investigation into the origin and composition of sICs predisposing to COVID-19 disease.

## Materials and Methods

### Subjects and specimens

Between March 2020 and April 2020, 41 patients with SARS-CoV-2 infection confirmed by real-time PCR were hospitalized in the University Medical Center, Freiburg. Serum samples were collected during hospitalization for routine laboratory testing. Clinical data were obtained from electronic medical records. A total of 27 patients necessitating invasive mechanical ventilation were included in the critical group. Fourteen patients requiring O_2_ supplementation were included in the severe group. Additionally, serum samples from 29 mild COVID-19 cases and 30 healthy donor (HD) plasma samples were used as controls in this study. For the SLE patient control cohort, sera were obtained from the Immunologic, Rheumatologic Biobank (IR-B) of the Department of Rheumatology and Clinical Immunology.

### Cell culture

African green monkey kidney Vero E6 cells (ATCC CRL-1586) were cultured at 37°C in Dulbecco’s Modified Eagle Medium (DMEM) supplemented with 10% (vol/vol) fetal calf serum (FCS, Biochrom), sodium pyruvate (1x, Gibco) and 100 U/ml penicillin-Streptomycin (Gibco). BW5147 mouse thymoma cells (BW, obtained from ATCC: TIB-47) were stably transduced with human FcγR as previously described (40, 41). Cells were maintained at 3×10^5^ to 9×10^5^ cells/ml in Roswell Park Memorial Institute medium (RPMI GlutaMAX, Gibco) supplemented with 10% (vol/vol) FCS, sodium pyruvate (1x, Gibco), 100 U/ml penicillin-Streptomycin (Gibco) β-mercaptoethanol (0.1 mM, Gibco). Cells were cultured at 37°C, 5% CO_2_. All cell lines were routinely tested for mycoplasma.

### Monitoring of antibody response to SARS-CoV-2 by ELISA

Serum IgG antibody titers targeting S1- and N-SARS-CoV-2 proteins were measured using commercial enzyme-linked immunosorbent assay (ELISA). Anti-S1-SARS-CoV-2 IgG was measured by the anti-SARS-CoV-2 ELISA (IgG) Euroimmune Kit (Euroimmune, Lübeck, Germany) according to manufacturer’s protocol. Results, expressed as arbitrary units (AU), were evaluated semi-quantitatively by calculation of the ratio of the extinction of the control or patient sample over the extinction of the calibrator. This ratio is interpreted as follows: < 0.8 negative; ≥ 0.8 to <1.0 borderline; ≥ 1.1 positive. Anti-N SARS-CoV-2 IgG was detected using the recomWell SARS-CoV-2 IgG Kit (Mikrogen Diagnostik GmbH, Neuried, Germany) according to manufacturer’s protocol. The corresponding antibody activity expressed in AU/ml is calculated using the formula (absorbance of sample / absorbance of cut-off) × 20. Results are interpreted as follow: < 20 negative; ≥ 20 to < 24 borderline; > 24 positive. IgG against the SARS-CoV-2 Spike Glycoprotein Receptor Binding Domain (RBD) were detected using SARS-CoV-2 IgG ELISA Reagent Set, kindly provided by InVivo (InVivo Biotech Services GmbH, Hennigsdorf, Germany) according to manufacturer’s protocol.

### Fcγ receptor activation assay

FcγRIIIA (CD16A, 158V) activation was measured by a cell-based assay as previously described (42). For detection of anti-S and anti-RBD-specific FcγR activation we utilized SARS-CoV-2-S- and RBD-coated plates (kindly provided by InVivo Biotech Services GmbH, Hennigsdorf, Germany). The recombinant (S)-protein was produced under serum-free conditions in mammalian cells and contains amino acid residues 1 to 1213 of the SARS-CoV-2 Wuhan-Hu-1-isolate (GenBank annotation QHD43416.1). The furin cleavage site was mutated, two mutations for protein stabilization were included, and the C-terminal domain was replaced by a T4 trimerization sequence and a C-terminal hexa-His-Tag (43). The recombinant RBD-protein represented amino acids 319 to 541 of the (S)-protein mentioned before. Both recombinant proteins were purified using immobilized metal exchange chromatography (IMAC) and preparative SEC under standard conditions in a regulated environment. Microtiter plates were coated using 0.2 μg recombinant (S)-protein or RBD-protein per well. N-specific FcγR activation was determined using plates coated with SARS-CoV-2-N (Mikrogen Diagnostik GmbH, Neuried, Germany). Respective plates were subsequently incubated with serial dilutions of SARS-CoV-2 positive sera or control sera in RPMI supplemented with 10% (vol/vol) FCS for 30 min at 37°C. All wells were thoroughly washed before co-cultivation with BW5147 reporter cells for 16 h at 37°C, 5% CO_2_. Cross-link activation of reporter cells was performed by direct coating of target antibody to ELISA plate (Nunc Maxisorp; 96 well, flat transparent), followed by a blocking step and incubation with 2 × 10^5^ reporter cells per well. For all activation assays, mouse IL-2 secretion was quantified by anti-IL-2 ELISA, as described earlier. FcγRIIIA (CD16A) activation by multimeric sICs was measured by a recently developed cell-based assay (25, 44). Briefly, 2×10^5^ BW5147-CD16 reporter cells were incubated with SARS-CoV-2 sera in a total volume of 200 μl for 16 h at 37°C, 5% CO_2_. Incubation was performed in a 96-well ELISA plate (Nunc Maxisorp) pre-treated with PBS containing10% FCS for 1 h at 4°C to avoid direct binding of serum IgG to the plate. Reporter cell mIL-2 secretion was quantified via ELISA as described previously (42).

### Purification of SARS-CoV2-S and –N specific antibodies from serum

SARS-CoV-2-specific antibodies were purified using SARS-CoV-2 spike protein (S)-coated plates (kindly provided by InVivo BioTech Services) and - nucleocapsid (N) - coated plates recomWell SARS-CoV-2 IgG (Mikrogen Diagnostik GmbH, Neuried, Germany). Patient sera were diluted 1:5 in 100 μl (two wells per serum sample) and incubated for one hour at 37°C with the S- and N-precoated plates. After washing using PBS-T (0.05% Tween 20) 100 mM formic acid (30 μl/well) was added and incubated for 5 min on an orbital shaker at room temperature (RT) to elute bound IgG. Following pH neutralization using TRIS buffer (1 M), the eluates were either directly processed or stored at 4°C.

### Quantitation of antigen-specific IgG amount

In order to determine the relative S1- and N-SARS-CoV-2 specific IgG antibody concentration of the generated eluates, S1- and N-ELISA were performed by the anti-SARS-CoV-2 ELISA (IgG) Euroimmune Kit (Euroimmune, Lübeck, Germany) and anti-N SARS-CoV-2 IgG ELISA (recomWell SARS-CoV-2 IgG Kit (Mikrogen Diagnostik GmbH, Neuried, Germany) as aformentioned.

### Analysis of antigen-specific IgG-Fc fucosylation

Fucosylation levels of S- and N-specific IgG were measured using a lectin-based ELISA assay. Briefly, 96-well Maxisorb plates (Nunc®) were coated with 50μl/well anti-human IgG-Fab fragment (MyBiosource, MBS674607) at a concentration of 2 μg/ml, diluted in PBS for one hour at 37°C. After three washing steps with PBS-T (0.05% Tween20) unspecific binding sites were blocked adding 300 μl/well Carbo-free™ blocking solution (VectorLab, Inc., SP-5040, LOT: ZF0415) for one hour at room temperature. After three further washing steps, eluted antibodies were serially diluted (2-fold) with PBS in a total volume of 30 μl/well and incubated for one hour at 37°C and 5% CO_2_. After washing (3x) using PBS-T, 50 μl/well of 4 μg/ml biotinylated Aleuria Aurantia lectin (AAL, lectin, VectorLab, B-1395) diluted in lectin buffer (10 mM HEPES, 0.1 mM CaCl_2_, 0.15 M NaCl, 0.1% Tween20) was added and incubated for 45 min at room temperature (RT). Following another three washing steps using PBS-T, Streptavidin-Peroxidase Polymer (Sigma, S 2438), at 1 μg/ml final concentration diluted in LowCross-HRP®-buffer (Candor, Order #.: 200 500) was added and incubated for one hour at RT. After washing five times with PBS-T, 50 μl/well of 1-Step™ Ultra TMB-ELISA Substrate Solution (ThermoFisher, 34028) was applied and the enzyme-substrate reaction was stopped after six minutes using 50 μl/well sulphuric acid (1 M H_2_SO_4_). Quantification of absorbance, OD_450nm_, was performed using a Tecan M2000. Relative fucosylation for each generated pool-eluate was calculated by normalizing OD_450nm_ (fucosylation) to its respective relative antigen-specific IgG amount.

### PEG Precipitation

Sera pools, consisting of eight different sera per pool, were diluted with varying amounts of PEG8000, in order to reach a final PEG8000 concentration of 1, 2, 3.5, 5 and 7.5% respectively. Mixtures were vortexed and incubated overnight at 4°C. For supernatant analysis, precipitates were sedimented via centrifugation at 13.000 rpm for 30 minutes at 4°C. For Mass Spectrometry analysis, PEG8000-precipitated sICs were shortly run into 10% polyacrylamide gels. After over-night fixation (40% ethanol, 10% acetic acid, 50% water) and washing (3x), complete lanes were excised.

### Benzonase treatment of sera

Serum from six individual patients containing CD16-reactive soluble immune complexes, were treated with 250 units (U) of Benzonase Nuclease (Sigma-Aldrich Chemie GmbH, Munich Germany) for 1 h at 4°C. After treatment, sera were titrated in complete BW5147 culture medium and tested for CD16 reactivity. Non-treated sera served as control. To verify Benzonase activity in the presence of human serum, 3 μg of pIRES-eGFP plasmid DNA (Addgene) were digested with 250 U of Benzonase. Successful nucleic acid digestion was visualized using a 1% agarose gel stained with Midori Green.

### Immune precipitation

For mass spectrometry analysis of SARS-CoV-2-S specific precipitates, individual sera containing CD16-reactive soluble immune complexes were subjected to immune precipitation (IP) using Pierce MS-compatible magnetic IP kit (ThermoFisher Scientific, Darmstadt, Germany) according to manufacturer’s protocol. Briefly 250 μl serum was incubated overnight at 4°C with 5 μg of biotinylated anti-RBD-specific TRES-1-224.2.19 mouse monoclonal antibody or TRES-II-480 (isotype control) (kind gift of H.M. Jäck, Erlangen) before addition of streptavidin magnetic beads. Beads were subsequently collected via centrifugation and elution buffer was added to detach putative precipitated antigen. The elution was dried in a speed vacuum concentrator and shortly run into 10% polyacrylamide gels. After over-night fixation (40% ethanol, 10% acetic acid, 50% water) and washing (3x), complete lanes were excised. Antibody biotinylation was performed using a Pierce antibody biotinylation Kit for IP (ThermoFisher Scientific, Darmstadt, Germany) according to manufacturer’s protocol.

### Mass Spectrometry

Proteins were in-gel digested with sequencing grade modified trypsin (Promega GmbH, Walldorf, Germany) similar to the procedure described by Pandey et al. (45). Vacuum-dried peptides were dissolved in 0.5% trifluoroacetic acid, loaded onto a trap column (C18 PepMap100, 5 μm particles, Thermo Fisher Scientific GmbH, Dreieich, Germany) with 0.05% trifluoroacetic acid (4 min, 10 μL/min) and separated on a C18 reversed phase column (SilicaTip™ emitter, 75 μm i.d., 8 μm tip, New Objective, Inc, Littleton, USA, manually packed 23 cm with ReproSil-Pur ODS-3, 3 μm particles, Dr. A. Maisch HPLC GmbH, Ammerbuch-Entringen, Germany; flow rate: 300 nL/min). For sample injection and multi-step gradient formation (eluent “A”: 0.5% acetic acid in water; eluent “B”: 0.5% acetic acid in 80% acetonitrile / 20% water; gradient length / acquisition time: 100 min or 175 min) an UltiMate 3000 RSLCnano system (Thermo Fisher Scientific GmbH, Dreieich, Germany) was used. Eluting peptides were electrosprayed at 2.3 kV via a Nanospray Flex ion source into a Q Exactive HF-X hybrid quadrupole-orbitrap mass spectrometer (both Thermo Fisher Scientific GmbH, Dreieich, Germany) and analyzed by data-dependent acquisition with HCD (higher energy collisional dissociation) fragmentation of doubly, triply and quadruply charged ions (loop count and dynamic exclusion dependent on the gradient length). Peak lists were generated with ProteoWizard msConvert (http://proteowizard.sourceforge.net/; version 3.0.11098), linear shift mass recalibrated (after a preliminary database search) using software developed in-house and searched against a database containing the SARS-CoV-2 UniProtKB reference proteome (proteome ID: UP000464024), all human UniProtKB/Swiss-Prot entries, and optionally (to reduce the number of incorrectly assigned matches) selected bacterial proteins (finally the Pseudomonas fluorescens (strain SBW25) reference proteome; proteome ID: UP000002332) with Mascot 2.6.2 (Matrix Science Ltd, London, UK; peptide mass tolerance: ± 5 ppm; fragment mass tolerance: ± 20 mmu; one missed trypsin cleavage and common variable modifications allowed).

### Neutralization assay

Serum neutralization capacity was analyzed as previously described (46). Briefly, VeroE6 cells were seeded in 12-well plates at a density of 2.8×10^5^ cells/well 24 h prior to infection. Serum samples were diluted at ratios of 1:16, 1:32 and 1:64 in 50 μL PBS total volume. Negative controls (PBS without serum) were included for each serum. Diluted sera and negative controls were subsequently mixed with 90 plaque forming units (PFU) of authentic SARS-CoV-2 (B.1) in 50 μl PBS (1600 PFU/mL) resulting in final sera dilution ratios of 1:32, 1:64, and 1:128. Following incubation at RT for 1 h, 400 μL PBS was added to each sample and the mixture was subsequently used to infect VeroE6 cells. After 1.5 h of incubation at RT, inoculum was removed and the cells were overlaid with 0.6% Oxoid-agar in DMEM, 20 mM HEPES (pH 7.4), 0.1% NaHCO_3_, 1% BSA and 0.01% DEAE-Dextran. Cells were fixed 48 h post-infection (4% formaldehyde for 30 minutes). Upon removal of the agar overlay, plaque neutralization was visualized using 1% crystal violet. PFU were counted manually. Plaques counted for serum-treated wells were compared to the average number of plaques in the untreated negative controls, which were set to 100%.

### Ethics

The protocol of this study conforms to the ethical guidelines of the 1975 Declaration of Helsinki and was approved by the institutional ethical committee of the University of Freiburg (EK 153/20). Written informed consent was obtained from participants and the study was conducted according to federal guidelines, local ethics committee regulations (Albert-Ludwigs-Universität, Freiburg, Germany: No. F-2020-09-03-160428 and no. 322/20; No 507/16 and 624/14 for the SLE patients).

### Statistical analyses

Statistical analyses were performed using linear statistical models. i.e. the two-group comparisons were made based on the t-statistic of the estimated effects. Differences over more than two groups were tested by Analysis of Variance (ANOVA) and multiple testing for subsequent two-group comparisons was then considered by performing Games-Howell post-hoc tests. For the time course data, patient differences were treated as random effects in a linear mixed effects model with time and clinical course (severe vs. critical) as fixed main and interaction effects. All analyses were performed at the log_2_ scale. Assumptions about variance heterogeneity and normal distribution were checked by visual inspection of diagnostic plots.

### Data and materials availability

All data associated with this study are present in the paper or Supplementary Materials.

## Results

### Patients and clinical information

We retrospectively analyzed serial serum samples collected for routine diagnostic testing from 41 patients hospitalized at our tertiary care center between March and June 2020 with SARS-CoV-2 infection confirmed by real-time PCR. Based on the clinical course, we categorized patients as either severely diseased (hospitalized with COVID-19 related pneumonia) versus critically diseased (COVID-19 related pneumonia and eventually in need of invasive mechanical ventilation). In total, 27 patients with critical and 14 with severe courses of disease were grouped into separate cohorts (Table 1). Most patients were older than 60 years with an overall mean age of 68 years (63 years and 76 years in the critically and severely diseased patients respectively). The majority of patients in both groups had comorbidities of different origin with cardiovascular diseases including hypertension representing the most frequent pathology (35/41, 85%). Similar to previous reports, high Interleukin 6 (IL-6) and C-reactive protein (CRP) levels were associated with severity of disease (Ø IL-6: 1452.1 pg/ml in the critical group vs 46.1 pg/ml in the severe group and Ø CRP: 162.2 mg/l vs 65.3 mg/l, Ø13-25 days post symptom onset respectively). Similarly, procalcitonin, a biomarker of microbial coinfection, was significantly higher in critically diseased patients (Ø value 9.9 ng/ml vs 0.17 ng/ml). Bacterial superinfection represented a further complication in 39% of the patients and was only slightly more frequent in patients with critical disease (11/27, 41% vs 5/14, 33%). More than half of the patients (59%) were treated with hydroxychloroquine/Lopinavir and Ritonavir (Kaletra®), (18/27, 67% in the critical group vs 6/14, 43% in the severe group). Notably, at the time of serum acquisition, only one patient received steroid treatment, which was given due to underlying chronic obstructive pulmonary disease. Finally, mortality rate was 37% (10/27) in critically and 7% (1/14) in severely diseased patients.

**Table 1:** Clinical characteristics of the hospitalized SARS-CoV-2 patients. Patients were categorized as either severely (hospitalized, requiring O_2_ supplementation, n=14) or critically diseased (hospitalized and in need of invasive mechanical ventilation, n=27). Diagnostic markers are depicted as mean and SD (in brackets) of all analyzed laboratory parameters obtained 13-25 days post symptom onset. Percentage [%] is indicated.

|  | All patients<br>n: 41 | % | critical<br>n: 27 | % | severe<br>n: 14 | % |
| --- | --- | --- | --- | --- | --- | --- |
| Age [years] | Ø 68<br>(31-90) | - | Ø 63<br>(39-79) | - | Ø 76<br>(31-90) | - |
| Female | 8 | 19.5 | 5 | 18.5 | 3 | 21.4 |
| Male | 33 | 80.5 | 22 | 81.5 | 11 | 78.6 |
| **Comorbidities** |  |  |  |  |  |  |
| Hypertension | 21 | 51.2 | 12 | 44.4 | 9 | 64.3 |
| Cardiovascular disease | 14 | 34.1 | 5 | 18.5 | 9 | 64.3 |
| Pulmonary disease | 6 | 14.6 | 2 | 7.4 | 4 | 28.6 |
| Chronic kidney disease | 6 | 14.6 | 1 | 3.7 | 5 | 35.7 |
| Diabetes | 10 | 24.4 | 6 | 22.2 | 4 | 28.6 |
| Malignancy | 8 | 19.5 | 4 | 14.8 | 4 | 28.6 |
| none | 6 | 14.6 | 6 | 22.2 | 0 | 0 |
| **Diagnostic markers** |  |  |  |  |  |  |
| Interleukin-6 [pg/ml] Ø | 1012.8 | - | 1452.1<br>(3774.6) | - | 46.1<br>(26.8) | - |
| Procalcitonin [ng/ml] Ø | 7 | - | 9.9<br>(21.9) | - | 0.17<br>(0.11) | - |
| C- reactive protein [mg/l] Ø | 128.1 | - | 162.2<br>(75.8) | - | 65.3<br>(47.1) | - |
| **Complications** |  |  |  |  |  |  |
| Bacterial superinfection | 16 | 39 | 11 | 40.7 | 5 | 35.7 |
| **Treatment** |  |  |  |  |  |  |
| Hydroxychloroquine, Ritonavir+<br>Lopinavir (Kaletra®) | 24 | 58.5 | 18 | 66.7 | 6 | 42.9 |
| **Fatal outcome** |  |  |  |  |  |  |
| Total | 11 | 26.8 | 10 | 37 | 1 | 7.1 |

### Kinetics of IgG antibody responses following symptom onset across severe and critical courses of disease

It has been observed that elevated SARS-CoV-2 antibody titers are associated with disease severity (15) and speculated to play a role not only in the clearance but also in the pathogenesis of SARS-CoV-2 infection (47). We initially analyzed the levels and kinetics of SARS-CoV-2 specific IgG in serial serum samples from patients hospitalized with critical (n=27) or severe (n=14) illness, a setting we also used in the following experiments. A total of 125 (critically diseased) and 79 (severely diseased) serum samples, obtained from the aforementioned patients at different time points within 6-25 days following symptom onset were analyzed by commercially available S1- and N- specific ELISA-based assays. Assay specificity was confirmed analyzing healthy donor (HD) serum samples (n=30) as negative control (Figure 1-figure supplement 1 A, B). Most patients developed detectable SARS-CoV-2 specific IgG responses within 9-14 days after symptom onset. SARS-CoV-2 specific IgG gradually increased over time in both severely and critically diseased patients reaching a plateau at 18-20 days after symptom onset (Figure 1 A, B). Varying antibody response kinetics were observed for each individual patient (Figure 1-figure supplement 2 A-D) with anti-N IgG titers rising significantly earlier than anti-S1 IgG (12.5 days ± 3.3 days vs 10.6 ± 3.8; p= 0.0091). A trend towards earlier seroconversion for anti-S1 IgG could be observed in critically diseased patients (mean time of seroconversion 11.4 ± 3.0 days in critically diseased patients vs 12.9 ± 3.8 days for severely diseased patients; p = 0.24), whereas time of seroconversion for anti-N IgG was similar in both groups (10.1 ± 3.2 and 10.4 ± 4.2 days for critically and severely diseased patients, respectively; p = 0.83). S1- and N-specific IgG levels at plateau did not significantly differ between the two groups. No significant difference between deceased and discharged patients was measured 13-25 days after symptom onset (Figure 1-figure supplement 1 C, D, E). Next, we evaluated and compared the neutralizing capacity of SARS-CoV-2 antibodies in either critically versus severely diseased patients in a plaque-reduction assay (Figure 1 C). All patients mounted a robust neutralizing antibody response (91% ± 10.5 % neutralization at a 1:64 serum dilution), with peaking titers at 18-20 days following symptom onset. Of note, two critically diseased patients developed a neutralizing response already at 6-8 days after symptom onset. In summary, we observed only minor differences in cohort wide kinetics of S1- or N-specific IgG levels between patients hospitalized with severe or critical clinical courses indicating that antibody levels per se did not correlate with severity of disease in our study.

**Figure 1.**
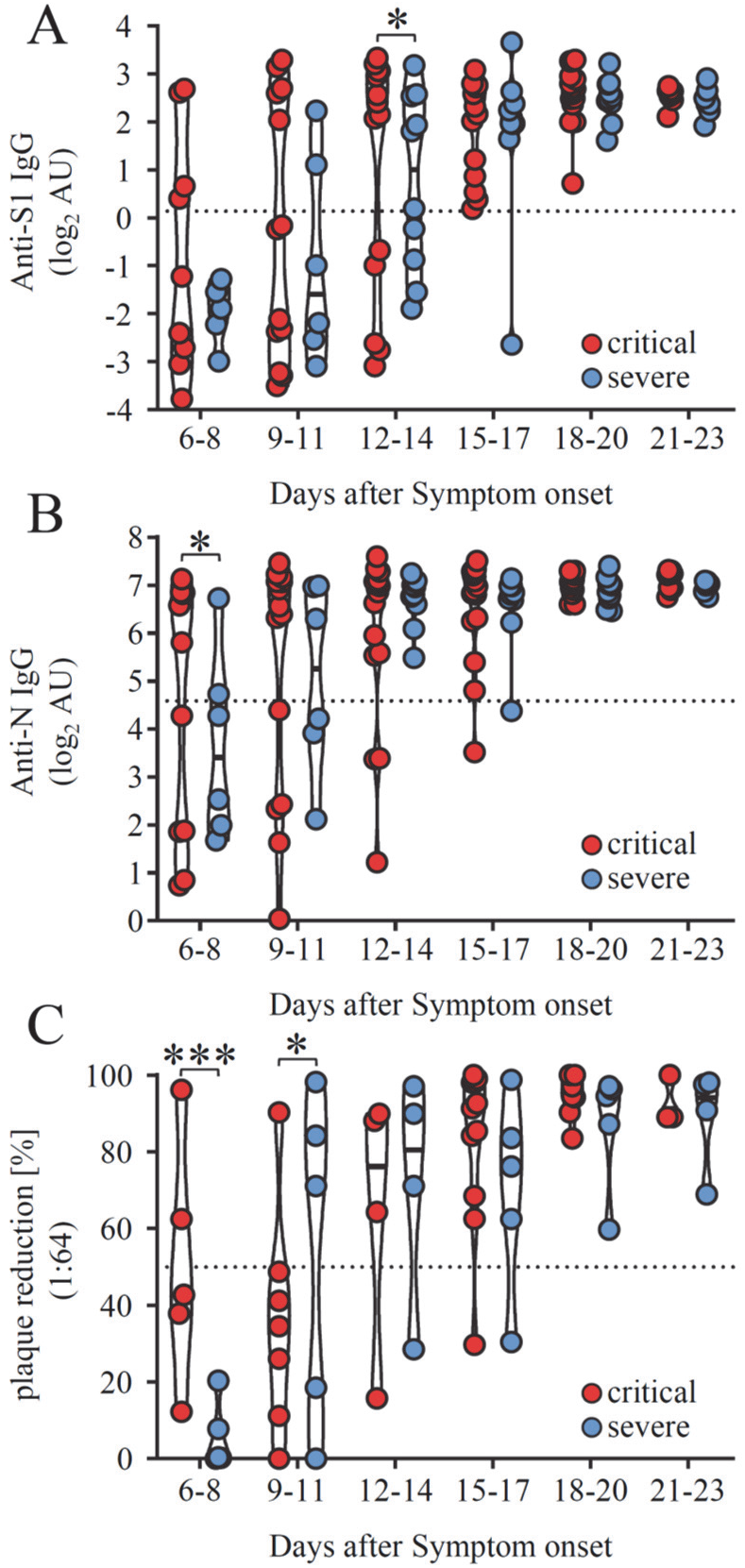
IgG responses against different SARS-CoV-2 proteins across severe and critical clinical course of disease. IgG antibody levels were analyzed in longitudinal serum samples from hospitalized SARS-CoV-2 infected individuals. 27 patients were categorized as critically diseased when in need of invasive mechanical ventilation (red symbols) compared to 14 severely diseased patients who did not require invasive ventilation (blue symbols). (A) IgG response against SARS-CoV-2 S1 –protein and (B) SARS-CoV-2 N-protein as determined by commercial ELISA assays. Dotted lines represent cut-off values for commercial S1- and N- specific ELISA assays. Each dot represents the mean value obtained by the analysis of all samples which were available at the indicated time points following symptom onset. Solid black lines indicate the median. (C) Serum neutralization capacity against SARS-CoV-2 measured by a plaque reduction assay. Sera were considered neutralizing upon 50% plaque reduction (dotted line) at a 1:64 dilution. Solid black lines indicate the median. Significant differences were tested using a linear mixed effects model (***, p<0.001; *, p<0.05).

### Patients with severe COVID 19 show enhanced FcγRIII/CD16 activation by S-specific IgG antibodies

FcγRIII (CD16) activation initiates multiple protective effector functions such as antibody-dependent cellular cytotoxicity (ADCC) by natural killer (NK) cells as well as antibody-dependent cytokine and chemokine secretion by NK cells and macrophages (18, 48). However, excessive FcγR stimulation can have severe adverse effects such as elevated cytokine release as observed in systemic autoimmune diseases or viral infections (18). Therefore, we hypothesized that an exaggerated FcγR mediated activation triggered by SARS-CoV-2 specific IgG might contribute to the exacerbation of COVID-19 in severely compared to critically diseased patients. To address this, we analyzed the ability of SARS-CoV-2 specific antibodies to activate CD16 (158V) using a previously validated cell-based reporter system (40-42, 49, 50) (Figure 2-figure supplement 1A). Considering the typically late time point of health deterioration, we performed an analysis of CD16 activation triggered by SARS-CoV-2 specific IgG with serum samples obtained 13-25 days following symptom onset (Figure 2). Sera were analyzed at a 1:500 dilution to stay within the dynamic range of detection (Figure 2-figure supplement 2). Depending on the availability of sample material 2-8 samples/patient/time-point were included in this analysis. If available in sufficient quantity, sera were reanalyzed. Reproducibility was tested using available serum surplus (Figure 2-figure supplement 3). Sera from 28 patients with mild SARS-CoV-2 infection and 30 healthy blood donors were included for reference. Semi-quantitative assessment of IgG titers using antigen-specific ELISA revealed comparable levels between critically and severely diseased patient cohorts (Figure 2 A, B, C). In contrast, S- (p=0.0147) and RBD-specific (p=0.0120) but not N-specific IgG-mediated CD16 activation was significantly increased in critically compared to severely diseased patients (Figure 2 D-F). Furthermore, normalizing CD16 activation to antigen-specific IgG titers, revealed significantly stronger CD16 activation by S- (p=0.0033) and N-specific (p=0.006) IgG compared to mildly diseased patients (Figure 2 G-I). Intriguingly, we observed a heterogeneous CD16 activation pattern characterized by either high or low CD16-activating sera irrespective of the clinical manifestation (Figure 2 D-F). Overall, a significant positive correlation could be determined between anti-SARS-CoV-2 antigen IgG titers and CD16 activation (Figure 2-figure supplement 4). Our data document a sustained CD16 activation by SARS-CoV-2 specific antibodies particularly in patients suffering from critical COVID-19 disease. Based on these results we confirmed the notion that elevated FcγRIII/CD16 activation by S- and or RBD-specific IgG might contribute to disease severity of COVID-19.

**Figure 2.**
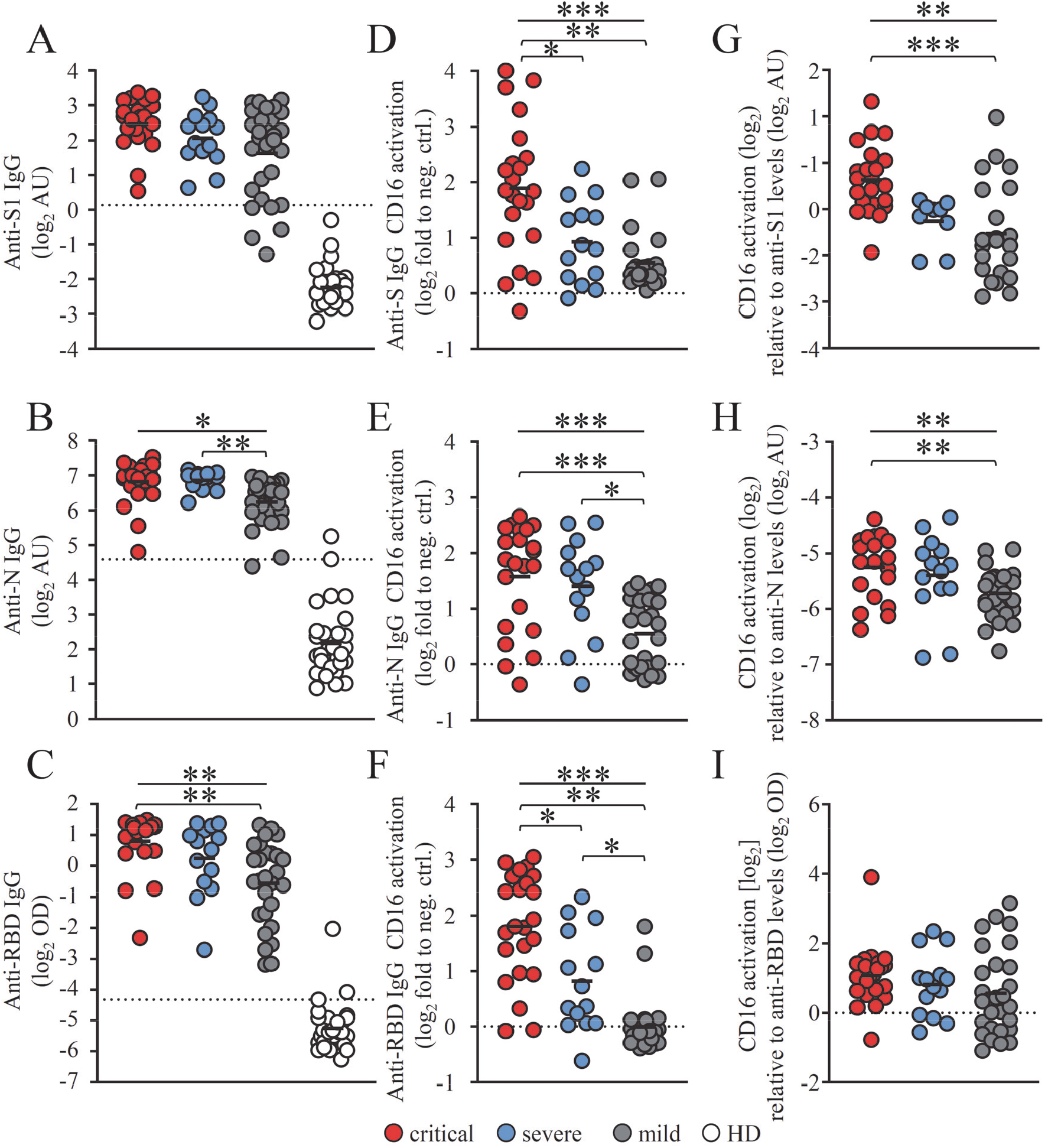
CD16 activation by SARS-CoV-2 - specific IgG is enhanced in critically diseased patients. FcγRIII activation by SARS-CoV-2-specific IgG on BW5147 reporter cells in serum samples obtained 13-25 days following symptom onset from 23 critically (red symbols) and 14 severely (blue symbols) diseased patients. Between 2 to 8 samples/patient were analyzed depending on the availability of sample material. Sera from 29 non-hospitalized patients with mild SARS-CoV-2 infection (grey symbols) and 30 healthy donors (open circles) served as reference. Each symbol represents the mean value of all available samples per patient. (A, B, C) ELISA levels for S1-N- and RBD-specific IgG. Dotted lines represent cut-off values for commercial S1-, N- and RBD - specific ELISA assays. Solid black lines indicate the mean. (D, E, F) FcγRIII activation by S-, N- and RBD-specific IgG expressed as log_2_ fold change relative to negative control. Solid black lines indicate the mean. (G, H, I) FcγRIII activation, expressed as log_2_ values relative to SARS-CoV-2-spcific IgG titers. Solid black lines indicate the mean. Significant differences over all three groups were tested by ANOVA and pairwise group comparison was made by Games-Howell post-hoc tests (***, p<0.001; **, p<0.01; *, p<0.05).

### Enhanced Fcγ-afucosylation of S-specific IgG in critically and severely diseased patients results in increased FcγRIII/CD16 activation

Based on the findings described above we speculated that differences in Fcγ mediated effector functions might contribute to disease severity of COVID-19. We compared CD16 high-versus CD16 low-activating patient sera regarding their SARS-CoV-2 specific IgG core fucosylation. Inspired by previous findings (51-53) we focused on determining IgG core fucosylation of S- and N-specific SARS-CoV2 IgG. To determine IgG core fucosylation we used a lectin-based ELISA preceded by antigen-specific antibody purification from immobilized SARS-CoV-2-antigen. Analysis of anti-S and anti-N IgG core fucosylation was performed on serum pools containing five sera of either critically or severely diseased patients obtained 13-25 days post symptom onset. Given the aforementioned heterogeneity in CD16-activation, we analyzed pools of 5 sera of either critically or severely diseased patients characterized by either high or low CD16-activation. To stay within the dynamic detection range, relative fucosylation was analyzed at a dilution of 1:4 (Figure 3). When analyzing serum pools from critically and severely diseased patients we determined a significantly lower level of core fucosylation among the high CD16 activators (Figure 3, plain-colored bars) compared to the low CD16 activators (Figure 3, shaded bars). This applied for both the S- and N-specific antibodies. These results are in line with previously published findings regarding the effect of Fcγ-afucosylation on FcγRIII/CD16 effector functions (51, 54) and recapitulate similar findings in the context of COVID-19 (52, 53). However, we did not observe significant differences between critically and severely diseased patients.

**Figure 3.**
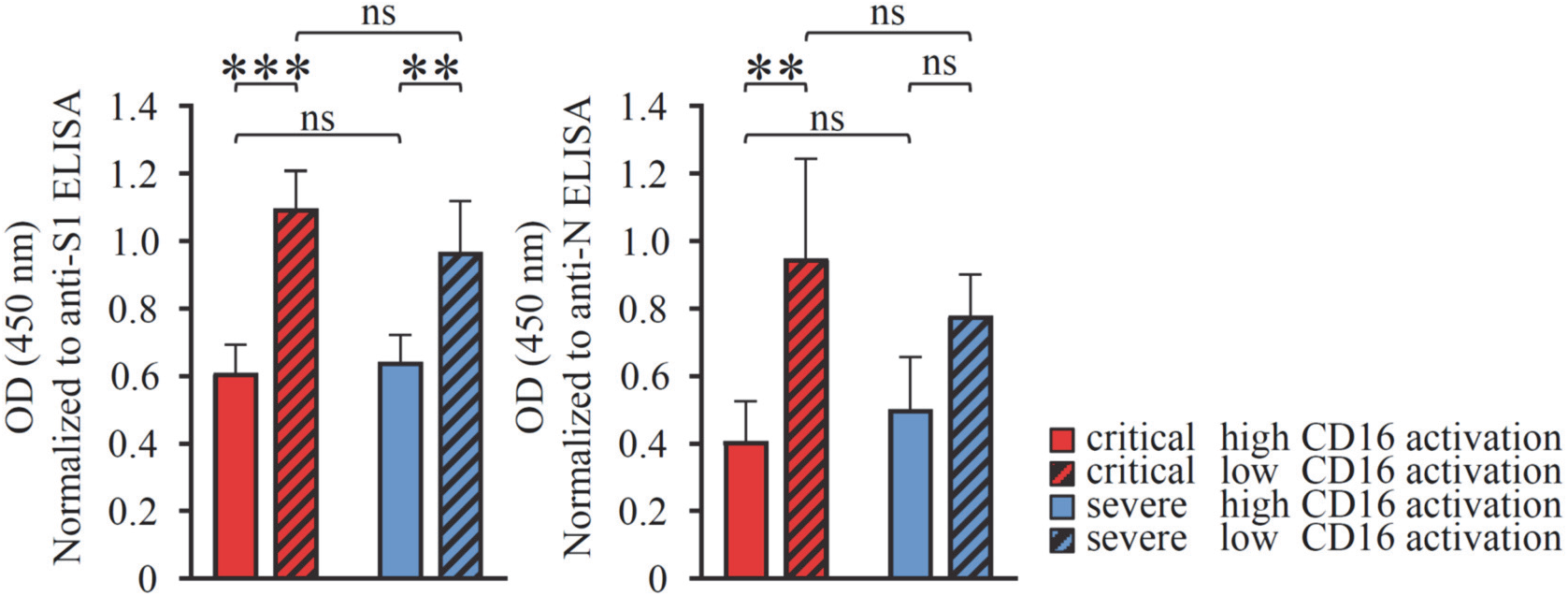
Anti SARS-CoV-2 IgG Fc core fucosylation in critical and severe COVID-19 cases. IgG-Fc core fucosylation levels of SARS-CoV-2 –specific IgG in critically (red bars) and severely (blue bars) diseased COVID-19 patients. Analysis was carried out on a pool of 5 different sera. Measured OD values for fucosylation of the generated eluates were normalized to their respective IgG titers determined by antigen-specific S1 and N ELISA. A) S-IgG-Fc-fucosylation and B) N-IgG-Fc-fucosylation in critically and severely diseased patients characterized by either high (red) or low (patterned) CD16-activation levels in the FcγR activation reporter assay. The mean and standard deviation (SD) of at least three independent experiments is depicted. Statistical tests using a two-factorial linear model indicate three significant differences between the low and high categories (***, p<0.001; **, p<0.01; *, p<0.05; ns = not significant).

### COVID-19 disease severity correlates with an increase in FcγRIII/CD16-reactive soluble IgG complexes

Next to afucosylation, it has been proposed that uncleared antigen-antibody immune complexes (ICs) might be involved in the pathogenesis of severe COVID-19.(34-36, 38). However, the actual presence of circulating, multimeric soluble ICs (sICs) in critically or severely diseased patients has not been shown yet. As extensive FcγR activation by sICs might contribute to the severe systemic inflammatory state occurring in some COVID-19 patients with prolonged disease, we surmised that sICs might be a putative explanation for the marked differences in IL-6, PCT and CRP levels between critically and severely diseased patients (Table 1). We thus set out to characterize our patient cohort regarding the presence of sICs in serum samples taken at various time points during disease and after hospitalization. To this end, we deployed a novel cell-based reporter assay developed to quantify CD16 (158V) activation by IgG-containing sICs, measuring their bioactivity (25, 44). This assay does not react to monomeric IgG or small dimeric complexes in solution, but specifically identifies multimeric sICs and has been successfully used to detect sICs in patients with systemic lupus erythematosus (SLE). In SLE, sICs are major driver of inflammation (24). This assay showed, as judged by conventional biomarkers, that sIC bioactivity correlates with SLE disease severity. Moreover, the assay is sensitive to sICs size with larger complexes leading to stronger receptor activation compared to small complexes (25). Analysis of serum samples, obtained 13-25 days after symptom onset, revealed the presence of highly CD16-reactive sICs in SARS-CoV-2 infected patients compared to healthy individuals (Figure 4A). Next, we compared sIC-mediated CD16 activation between COVID-19 patients of varying disease severity. While all COVID-19 patient groups tested positive for reactive sICs compared to healthy control (HD) sera, we found that critically diseased patients show a striking increase in reactive sICs compared to patients with severe or mild disease (Figure 4B). We then compared sIC bioactivity between sera from critically diseased COVID-19 patients and sera from SLE patients with active disease (Figure 4C). We conclude that sICs formed in COVID-19 are comparable to sICs formed during active SLE regarding their potential to drive inflammation. Only 6 out of 27 patients with critical disease (22%) showed no sIC-mediated CD16 activation. As we did not detect highly reactive sICs in the serum of 47 patients with acute respiratory distress syndrome (ARDS; mean age 57.5 years) in response to infections of different etiology including CMV reactivation, HIV/AIDS, influenza or pulmonary TBC infection, we conclude that the formation of reactive sICs is associated with severe SARS-CoV-2 disease (Figure 4-figure supplement 1). Remarkably, longitudinal analysis of reactive sICs in the serum of critically or severely diseased patients revealed high CD16 activation levels in 4 critically diseased patients already 6 to 8 days after symptom onset (Figure 4D). Of note, 2 of 4 patients with an early increase of circulating reactive sIC eventually died. sIC-mediated CD16 activation persisted in 14 of 19 critically diseased patients at high levels until day 26 after symptom onset. sIC-mediated CD16 activation in severely diseased patients was slightly delayed compared to critically diseased patients and was first detected in 4 patients 9-11 days after symptom onset (Figure 4D). Only 4 of 14 patients with severe disease showed detectable sIC-mediated CD16 activation. To verify that sICs represent the CD16-reactive component in the serum of COVID-19 patients, we analyzed serum-mediated CD16 activation before and after PEG8000-precipitation. This treatment was previously shown to selectively precipitate large IgG complexes from solution (25, 55). For this analysis, pools of 8 sera, showing either high (IC+) or no (IC-) CD16 activation, were compared. Sera from healthy donors (HD) served as a negative control. Compatible with the hypothesis of serum-derived sICs driving CD16 activation, no activation was observed following incubation with 3.5% PEG8000 (Figure 4-figure supplement 2 A). To ensure that the treatment did not precipitate monomeric IgG, we tested the depleted sera for remaining S1- and N-specific IgG. As depicted S1- and N- specific IgG could still be detected at unchanged high levels in samples treated with 3.5% PEG8000 (Figure 4-figure supplement 2 B). When resolving sIC-mediated CD16 activation over the complete time of hospitalization for select patients from which samples at different time points were available, we observed that sIC reactivity predominantly precedes anti-S1 IgG in ELISA as well as CD16 activation by SARS-CoV-2-specific IgG (Fig. 4-figure supplement 3, Fig. 4-figure supplement 5). This implies that sIC formation does not depend on the presence of SARS-CoV-2 antigens. Accordingly, we were not able to identify any SARS-CoV-2-derived antigens in PEG8000-precipitated sICs using tandem mass spectrometry (data not shown). To further exclude the formation of multimeric sICs formed from circulating S1 antigen, we also specifically targeted S1 for precipitation from patient serum using biotinylated S1-specific monoclonal antibodies. However and in line with our previous approach, S1-specific precipitation using streptavidin-sepharose beads and subsequent mass spectrometry analysis for any SARS-CoV-2-specific antigens in sICs remained without result (data not shown). Recently, the role of neutrophil mediated intravascular NETosis was reported to play a critical role in thrombose formation and subsequent organ damage observed in severe clinical forms of COVID-19 (56-58). Since this process could mediate the formation of aggregated IgG as a form of sICs, we next tested whether Benzonase® nuclease treatment of patient serum would dissolve reactive sICs. To this end we tested sera from critically diseased patients or healthy individuals and compared CD16 reactivity before and after nuclease treatment (Figure 4-figure supplement 4). Nuclease activity in diluted human serum was controlled using plasmid DNA for reference. This revealed that nucleic acid was not involved in the formation of CD16-reactive sICs in critically diseased patients. Finally, we tested pooled patient sera for autoantibodies against a panel of prototypical autoantigens associated with autoimmune disease including anti-nuclear autoantibodies (ANA) by indirect immunofluorescence, dsDNA autoantibodies by ELISA and autoantibodies against the extractable nuclear antigens (nRNP/Sm, Sm, SS-A, Ro-52, SS-B, Scl-70, PM-Scl, Jo-1, CENP B, PCNA, nucleosomes, histones, ribosomal P-protein, AMA-M2, DFS70) by dot blot in case SARS-CoV-2 infection triggers autoantibody formation and possible sIC formation. However, no significant autoantibody titers could be detected in any sera pool (data not shown). Although we were not able to identify their origin, our data clearly indicates the presence of circulating sICs in COVID-19 patients with an increase in CD16-reactive sICs corresponding with severity of disease and reaching activation levels comparable to those observed in SLE patients with active disease. Accordingly, we conclude that circulating sICs are a hitherto unknown, yet contributing factor to COVID-19 disease severity and, regarding infectious diseases, our findings represent an observation unique to severely diseased COVID-19 patients.

**Figure 4.**
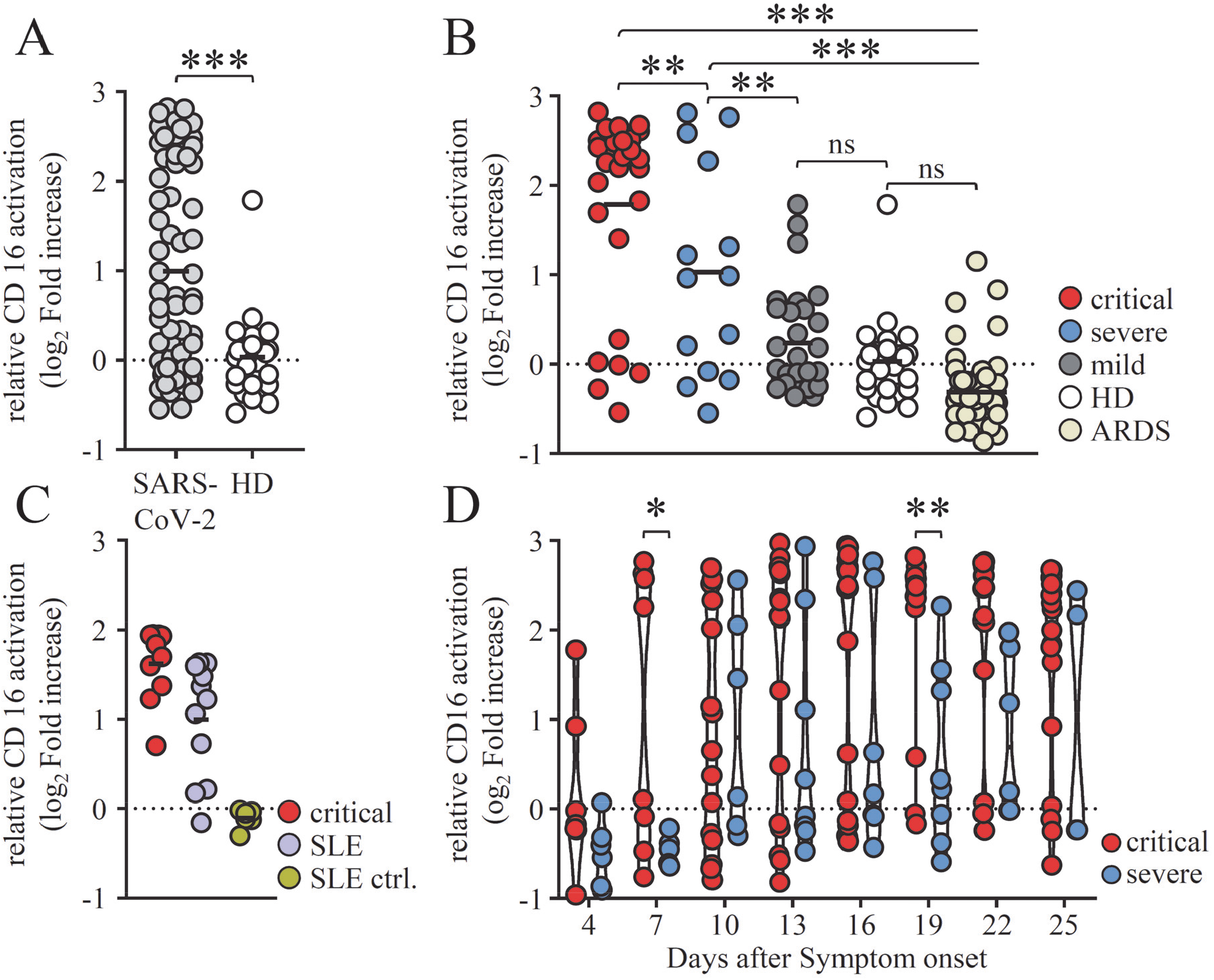
Severe COVID-19 disease coincides with high CD16 activation by sICs. Serial serum samples obtained 13-25 days after onset of symptoms were analyzed in a cell-based reporter assay which is sensitive to sIC amount and size (25, 44). FcγR activation is shown as log_2_ fold change relative to negative control. Each symbol represents the mean value obtained by the analysis of all samples available in the indicated time range for each individual patient. A) Analysis of CD16 activation by sICs in SARS-CoV-2-infected patients compared to healthy blood donors B) Levels of IC-mediated CD16 activation across severe, critical and mild clinical courses of COVID-19 disease, in healthy donors (HD) and in non-COVID-19 patients who developed acute respiratory distress syndrome (ARDS). Solid black lines indicate the mean. Two-group comparisons with the linear model indicate significant differences between critical cases and all other groups, as well as between severe cases and all other groups (***, p<0.001; **, p<0.01). No significant differences (p>0.05) have been found for the comparisons mild vs. healthy and for HD vs. ARDS. C) Select sera from critically diseased patients were compared to sera from SLE patients with active disease regarding CD16 activation. Sera from healthy donors served as SLE-negative control. Solid black lines indicate the mean. D) Kinetics of IC-mediated CD16 activation in critically and severely diseased patients. Days after symptom onset are depicted as a range (+/- 1 day). Solid black lines indicate the median. The mixed effects model indicates two time points with significant differences (**, p<0.01; *, p<0.05).

## Discussion

We collected and analyzed data from 41 COVID-19 patients hospitalized at the University Hospital Freiburg. Patients were categorized by severity of disease into severely (n=14) and critically diseased patients (n=27). Both groups were of comparable average age and had a similar male-to-female ratio. For comparison we also analyzed 28 mildly diseased and 30 healthy individuals. As key findings we identify *de novo* produced afucosylated SARS-CoV-2 IgG and the presence of soluble circulating immune complexes (sICs) activating FcγRIII/CD16 as potential risk factors closely associated COVID-19 severity.

### Circulating sICs contribute to COVID-19 disease severity

Using an adapted reporter cell activation assay optimized to detect sICs (25, 44) we provide first evidence of circulating sICs in the serum of COVID-19 patients and experimentally confirm previous hypotheses suggesting immune complexes as potential drivers of disease progression in COVID-19 (34-36, 38). In fundamental contrast to opsonized antigens decorating virus-infected cells in tissues, sICs become distributed systemically. Thus constitutive activation of CD16^+^ monocytes, granulocytes and NK cells could readily explain systemic responses which potentiate local inflammation in virus-infected tissues intensifying organ damage and dysfunction. Although the origin of the circulating immune complexes and the nature of the bound antigens still remains elusive, we clearly show that the presence of IgG-containing sICs during SARS-CoV-2 infection is directly responsible and sufficient for the observed FcγRIII/CD16 activation by patient serum. Recent work has shown that viral antigens can be detected in the serum of patients (59, 60). However, as we find sIC reactivity to predominantly precede SARS-CoV-2-S specific IgG responses, we conclude that circulating S or shed S1-antigens are not involved in sIC formation. Since sICs are commonly associated with immunopathology in autoimmunity (23, 24, 61) and several studies have described that a variety of specific auto-antibodies can be detected in certain critically ill COVID-19 patients (62-64), we could not identify a distinct culprit antigen linked to sIC formation when searching for prototypical autoantibodies. However, as sIC formation is strongly reminiscent to SLE and sICs initiate a common terminal pathway of inflammation we classified patients as sIC-prone or non-sIC-prone (graphical abstract). Of note, besides sIC formation a range of additional phenotypical abnormalities shared between B cell populations in autoimmune disorders exemplified by active SLE and severe COVID-19 have been observed. This includes the pronounced engagement of extrafollicular B cell responses, associated with the activation of effector B cells lacking naïve (IgD) and memory markers (CD27) as well as class-switched antibody secreting cells (62).

We suggest a hidden predisposition in sIC-prone patients resulting in a strong early inflammatory response to SARS-CoV-2 infection possibly being a trigger for further sIC formation and the generation of afucosylated SARS-CoV-2 IgG. Very recent work reports that an acute SARS-CoV-2 infection triggers the *de novo* IgG production against multiple autoantigens. In this study, 60-80% of all hospitalized COVID-19 patients exhibited anti-cytokine IgG (ACA) (65). The authors show that ACA levels and specificity change over time during hospitalization, suggesting ACA induction in response to viral infection and inflammation. Further, it has been shown that pre-existing neutralizing anti-type I interferon antibodies, which can be found in about 10% of patients with severe COVID-19 pneumonia, are related to the highest risk of developing life-threatening COVID-19 disease (66). Therefore, the *de novo* induction of anti-cytokine auto-antibodies in a large proportion of hospitalized COVID-19 patients as described by Chang et al. (65), might indeed represent a source of circulating sICs in COVID-19. In such a scenario, immune responses are deviated first by an immunodepletion of critical cytokines and second through the formation of pathological sICs which trigger immunological damage. We show that critically diseased patients exhibit significantly higher levels of reactive sICs compared to less severely diseased patients. Notably, CD16 activation levels in patients with critical disease were comparable to those measured in SLE patients, where circulating sICs have long been shown to crucially contribute to tissue damage and disease manifestations (67, 68). In addition, sIC responses can be found significantly earlier in critically diseased patients, which was associated with a fatal disease outcome. We also find that patients show a wide range of sIC reactivity. According to the Heidelberger-Kendall precipitation curve (69), sIC size is critically dependent on the antigen:antibody stoichiometry. As the used FcγR activation assay is highly sensitive to sIC size (25), it is likely that this also plays a role when detecting this bioactivity in COVID-19 patient serum. Therefore, we propose that in addition to the presence of sICs, the size of sICs plays a role in CD16 driven COVID-19 immunopathology. Based on these findings, together with the higher levels of afucosylation, we conclude that CD16 activation in COVID-19 disease is governed by sIC formation and IgG glycan profiles (Figure 5). It can be hypothesized that the formation of sICs in predisposed patients initiates a vicious circle of FcγR-mediated inflammation leading to an increase of IgG afucosylation, followed by enhanced FcγR activation by SARS-CoV-2-specific IgG, further contributing to inflammation and, conceivably, to *de-novo* sIC formation. Indeed, there is evidence in this direction from a clinical perspective provided by a recent study that finds the administration of intravenous immunoglobulin (IVIg) to alleviate COVID-19 disease (70). Although no direct proof, this heavily implies that the saturation of FcγRs mitigates immunopathology as previously reported for autoimmune diseases (71). Therefore, our findings provide an explanation for the sustained immunopathology following SARS-CoV-2 infection observed in some patients as well as for the efficacy of IVIg treatment in severe to critical COVID-19 disease. Finally, when we analyzed sera from COVID-19 patients obtained during the following waves of the pandemic, we again found comparable levels of reactive sICs in the serum of critically diseased patients (data not shown) implying that sIC formation in COVID-19 is conserved across different SARS-CoV-2 strains. It will be important to investigate whether such sICs may persist in reconvalescent patients and might be an explanation for immune alterations including auto-antibodies observed in patients with persistent long COVID-19 symptoms (72).

**Figure 5.**
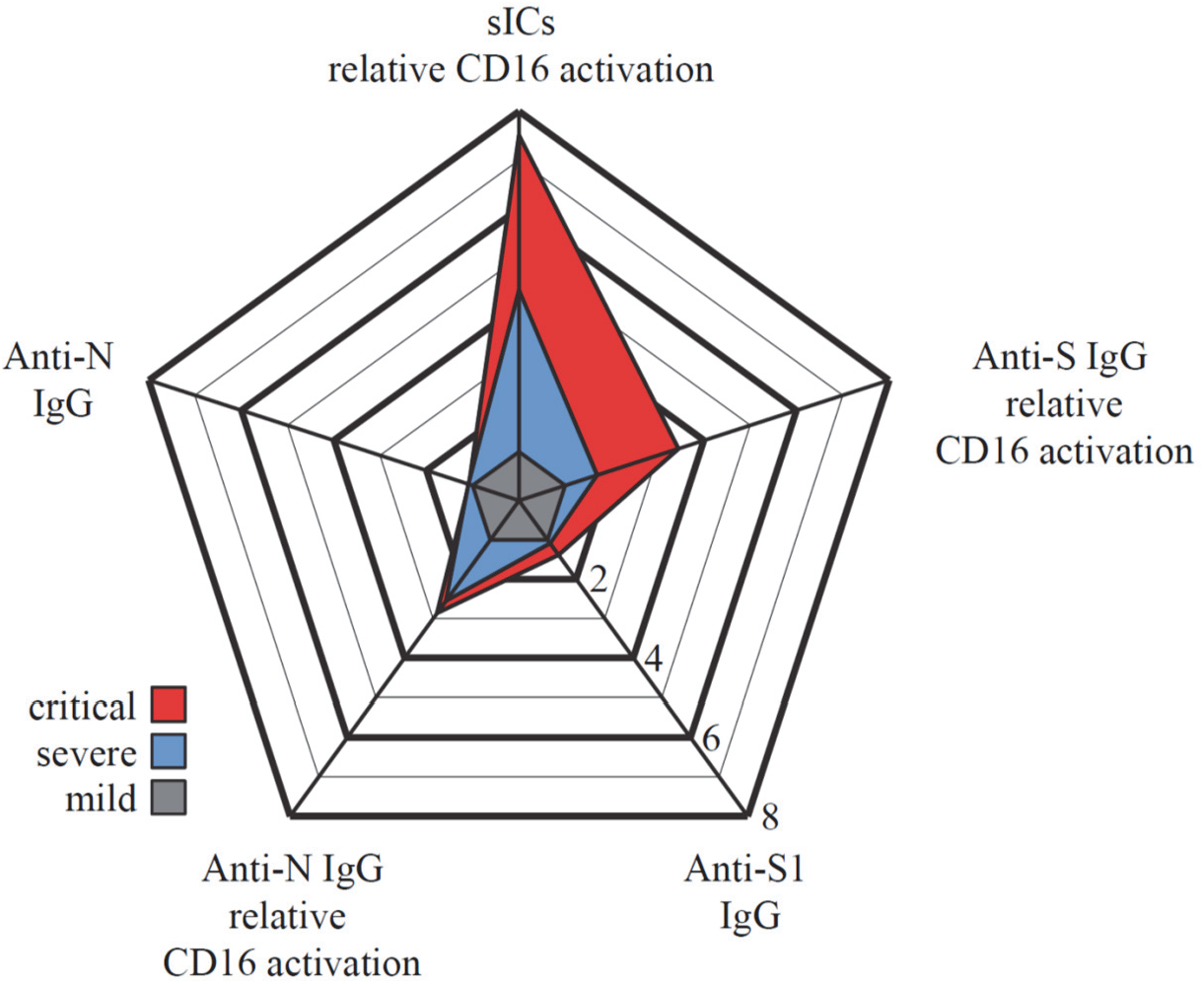
Summary of antibody features from SARS-CoV-2-infected patients with critical and severe disease. Relative multivariate antibody features illustrated as radar chart in critically (red) or severely (blue) diseased COVID-19 patients normalized to the corresponding features of patients with mild infection (grey). Each spoke represents one of the following variables: ELISA (S1-IgG, N-IgG,) and CD16 activation (S-IgG, N-IgG, multimeric sICs). Arithmetic mean values of log_2_ values were calculated for each group (days 13-25 post symptom onset) respectively. The fold change compared to mildly diseased patients is shown.

## Acknowledgements

We thank Sophia Ruben and Torsten Schulz from InVivo Bio Tech Services for preparing SARS-CoV-2 (S)- and RBD-coated plates and preparing the SARS-CoV-2 IgG ELISA Reagent Set. We thank Hans-Martin Jäck for providing TRES-1-224.2.19 and TRES-II-480 monoclonal antibodies and Quinnlan David for critically reading the manuscript. We also thank Andreas Schlosser for validation of our Mass Spectrometry data.

## Author contribution

Conceived and designed the experiments: J.A., A.M-P., S.G., P.K., A.L, V.F., M.S., H.H.

Performed the experiments: J.A., N.G., U.S., S.G., K. C., W.B.

Analyzed the data: J.A., S.G., P.K., V.F., K.C., A.M-P., W.B., C.K.

Contributed/reagent/sample material: A.B.G., D.H., T.W., NG.M., RE.V.

Writing and original draft preparation: J.A., S.G., P.K., V.F.

Review and editing: H.H., M.S., K.C.

Conceptualization: V.F., H.H.

## Supplementary Figures

**Figure 1-figure supplement 1.**
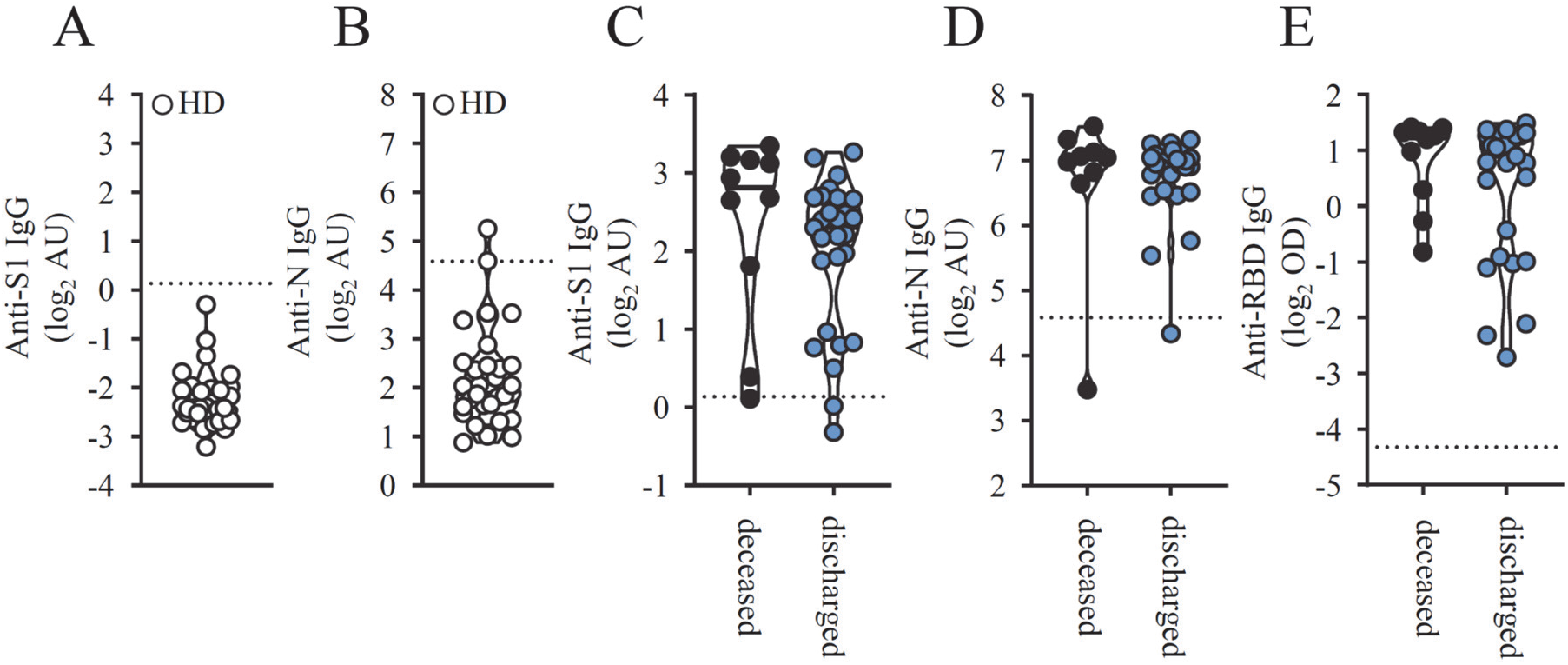
SARS-CoV-2 specific IgG levels in seronegative patients and according to disease outcome. A) S1- and B) N-specific IgG levels in 30 healthy donors. Solid black lines indicate the median. C) Cumulative S1-, D) N- and E) RBD-specific IgG levels measured 13-25 days after symptom onset in deceased (black symbols) and not deceased COVID-19 patients (blue symbols). Each symbol represents the mean value obtained by the analysis of all samples available in the indicated time range for each individual patient. Solid black lines indicate the median.

**Figure 1-figure supplement 2.**
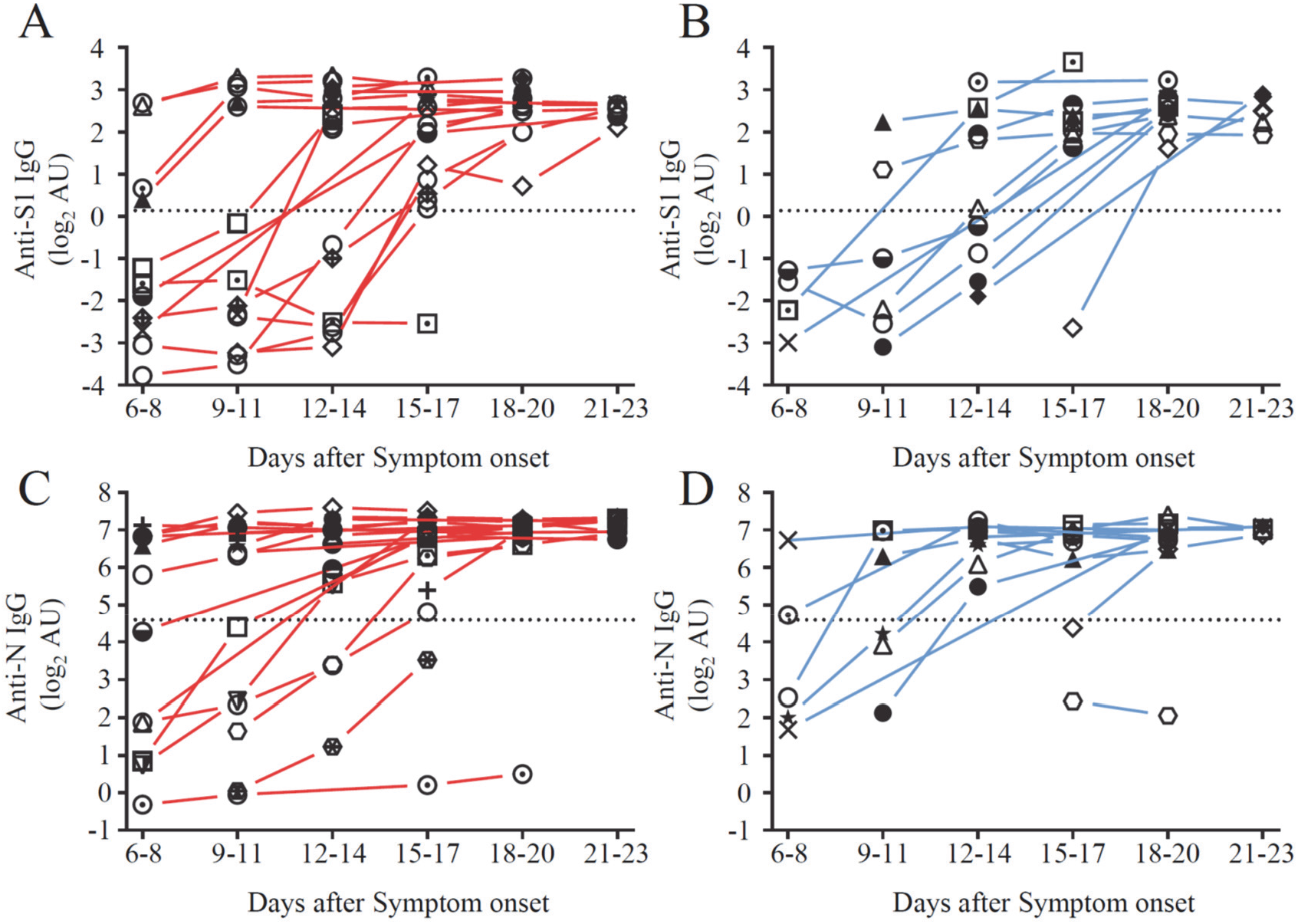
Longitudinal changes in anti- SARS-CoV-2 IgG titers in severely and critically diseased patients. Serial serum samples were collected from hospitalized COVID-19 patients and used for SARS-CoV-2–specific IgG measurement. IgG responses against SARS-CoV-2 S1- and N-protein in (A, C) critically (red symbols) and (B, D) severely (blue symbols) diseased patients. Dotted lines represent cut-off values for commercial S1- and N- specific ELISA assays. Each symbol represents the mean value of all samples which were available for each patient at the indicated time range after symptom onset. There are no significant t-tests (i.e. p>0.05 for all comparisons).

**Figure 2-figure supplement 1.**
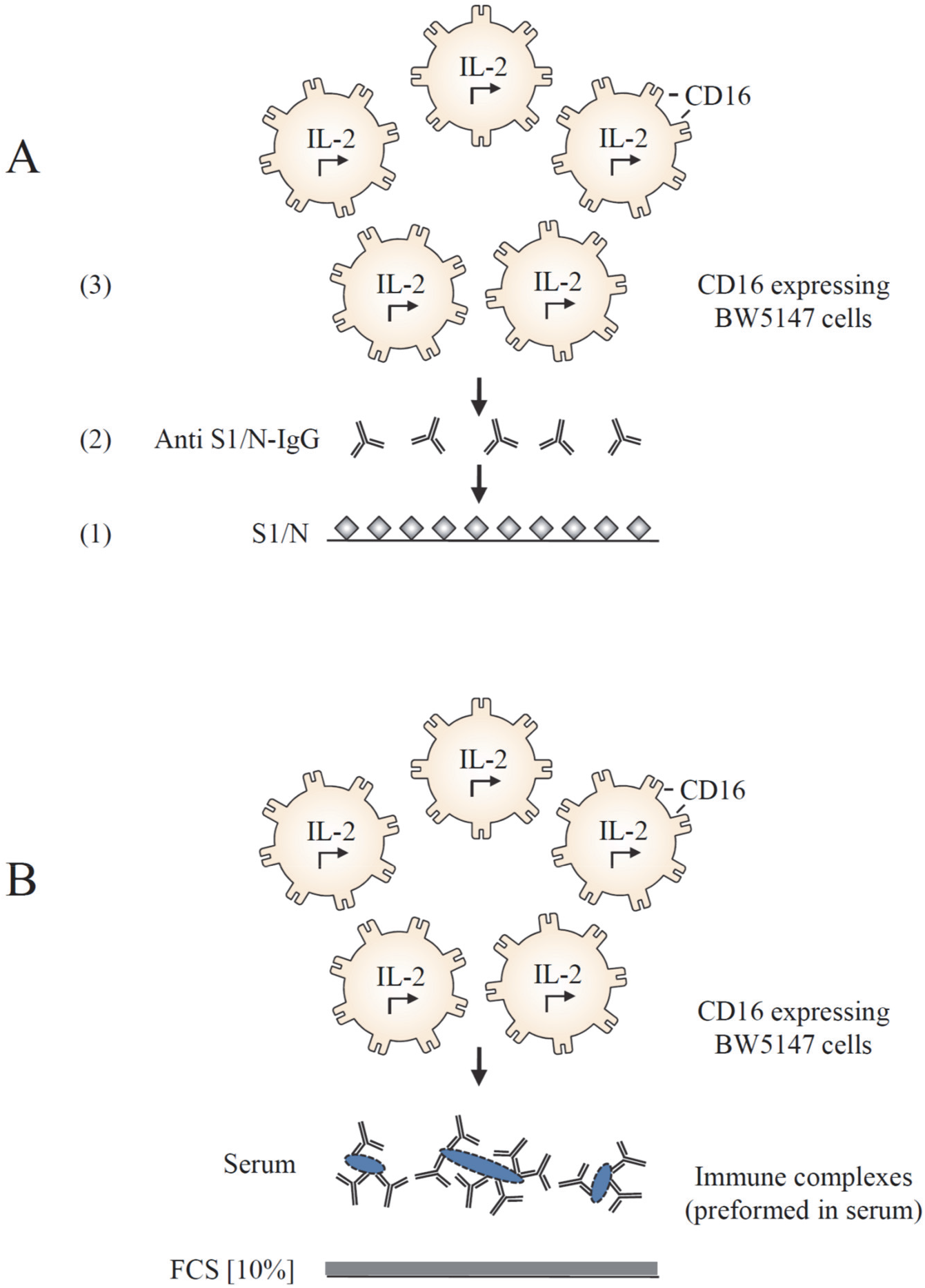
Cell-based reporter assay measuring CD16 activation in response to immobilized IgG and sICs. BW5147 reporter cells expressing chimeric human FcγRIII secrete IL-2 in response to FcγR activation by A) clustered viral specific IgG binding solid-phase antigen or B) soluble ICs. Solubility of sICs is achieved by pre-blocking an ELISA plate with PBS supplemented with 10% FCS as previously described (25, 44).

**Figure 2-figure supplement 2.**
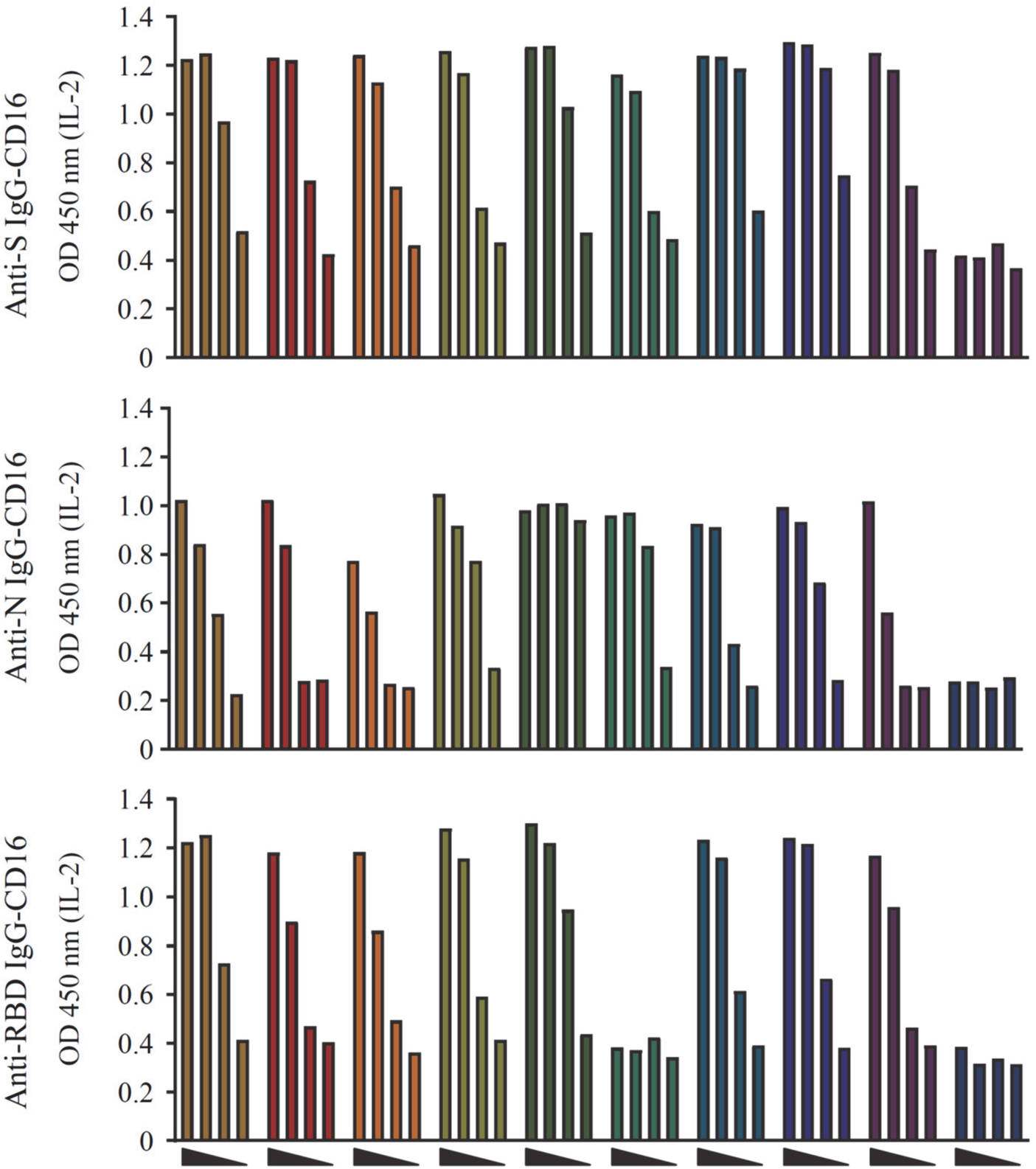
Dose dependent CD16 activation by SARS-CoV-2 specific IgG. CD16 activation by A) S-, B) N- and C) RBD-specific IgG in 9 representatively selected serum samples and one SARS-CoV-2 negative serum (dark blue bars). Sera were serially diluted at 1:20, 1:100, 1:500 and 1:2500. FcγRIII activation initiates IL-2 secretion by reporter cells, which is subsequently measured via ELISA (OD 450 nm). Based on this empirical pretesting all sera were thereafter tested at 1:100 and 1:500 dilutions to reach an optimal dynamic range of response. The OD values obtained by the 1:500 dilutions were used for subsequent data analysis.

**Figure 2-figure supplement 3.**
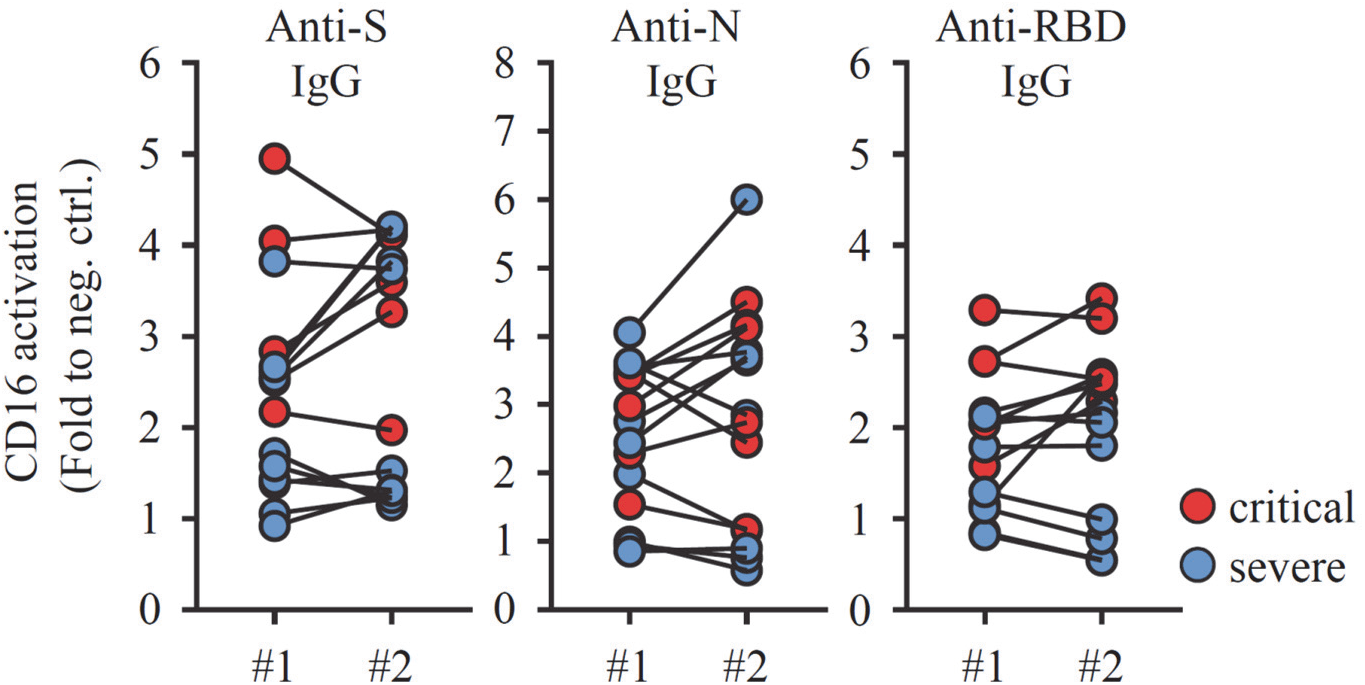
Reproducibility of CD16 activation measurements by SARS-CoV-2 specific IgG. Selected sera which were available in sufficient amount from patients with critical (red symbols) or severe (blue symbols) SARS-CoV-2 infection were tested in two independent experiments to show reproducibility and consistency of results. CD16 activation by S-, N- and RBD specific IgG is shown. Statistical tests using a Kolmogorov-Smirnov test indicate no significant differences.

**Figure 2-figure supplement 4.**
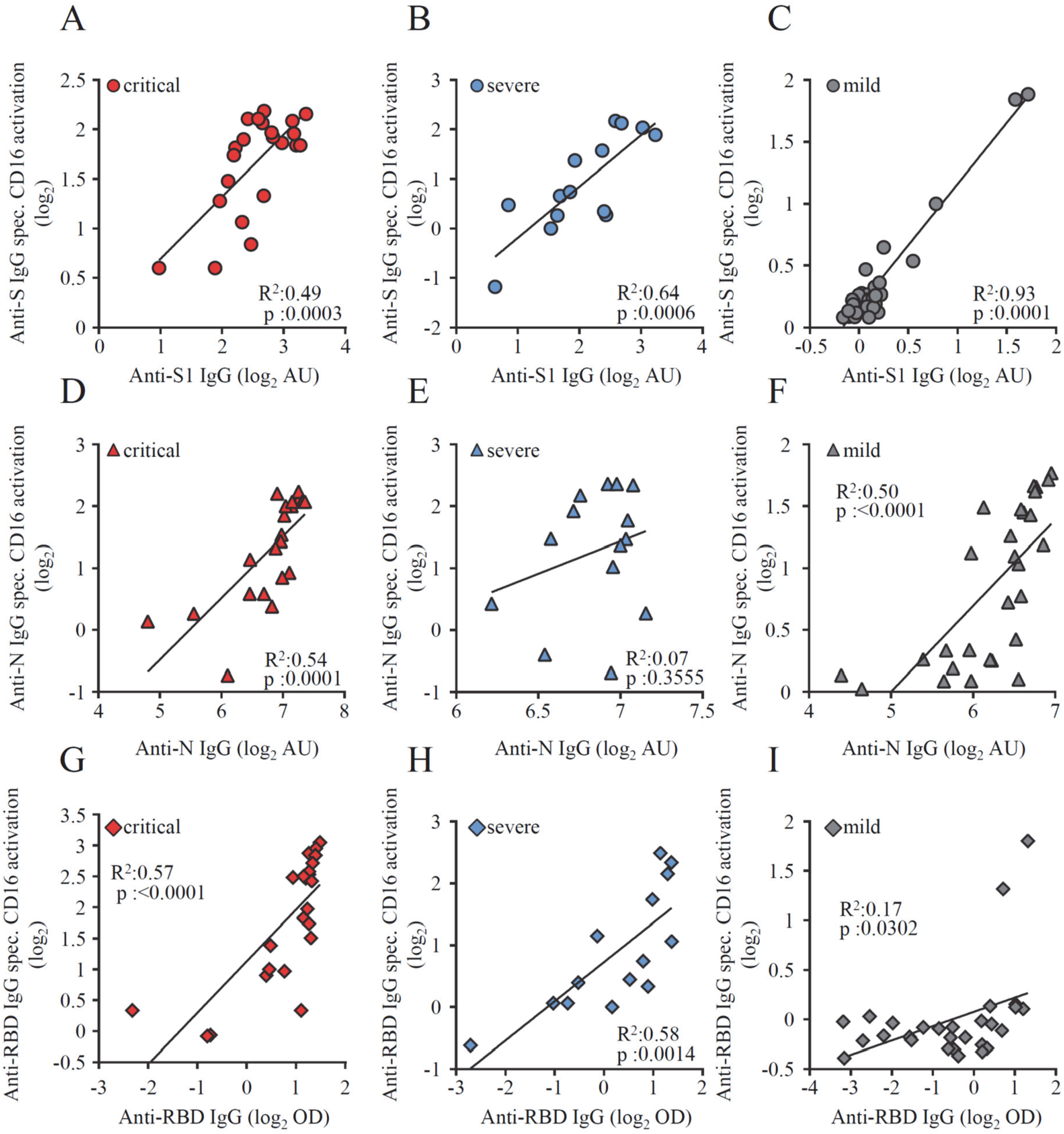
Correlation of CD16 activation by virus specific IgG and ELISA levels. Pearson’s correlation coefficient was used to assess the relationship between virus–specific IgG levels and their capability to trigger CD16 activation on BW5147 reporter cells in 22 paired samples from patients with critical disease (red symbols), 14 paired samples from patients with severe disease (blue symbols) and 28 samples from patients with mild disease (grey symbols). Each dot represents the mean value obtained by the analysis of all samples available at the indicated time points. (A-C) anti-S IgG, (D-F) anti-N IgG and anti-RBD-IgG (E-I).

**Figure 4-figure supplement 1.**
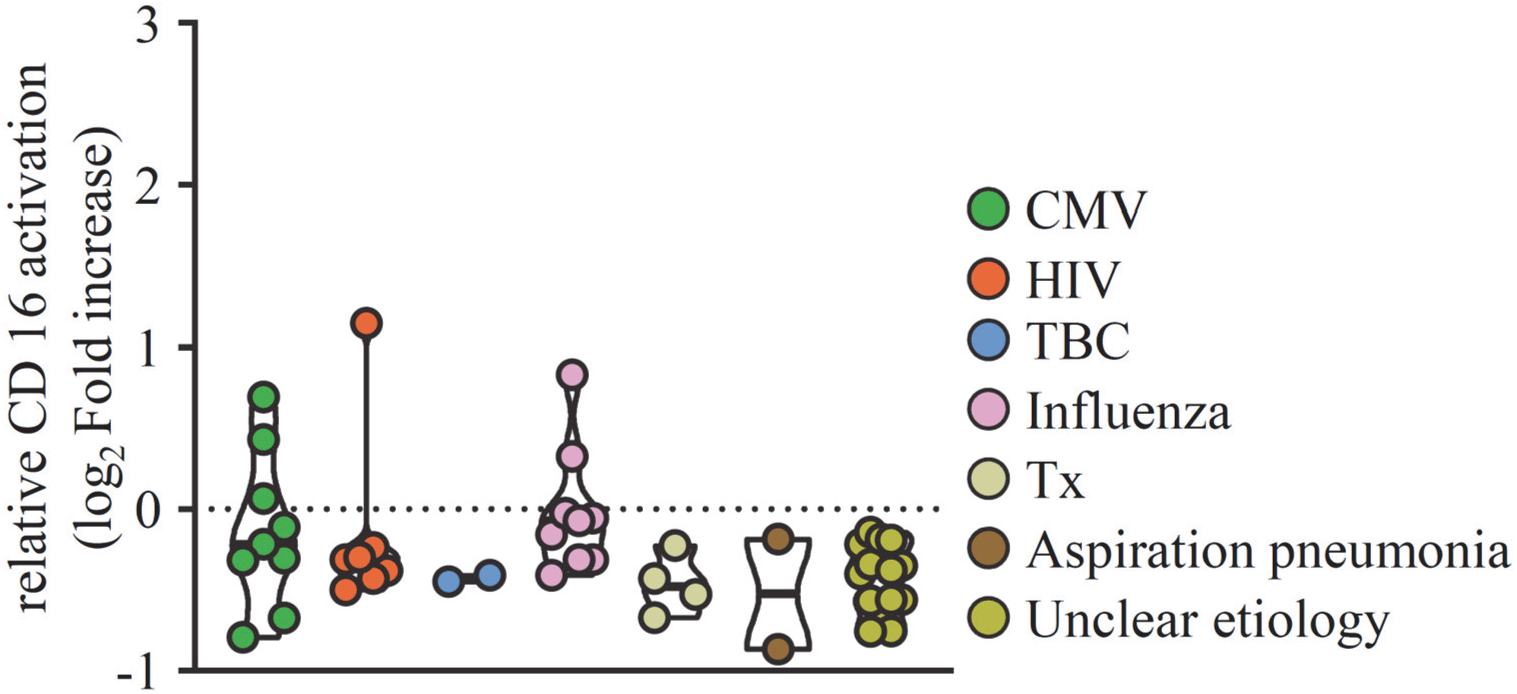
CD16 activation by sICs in non-COVID-19 patients with ARDS. Serum samples from 47 patients with ARDS in response to infections of different etiology were analyzed in a cell-based reporter assay which is sensitive to sIC amount and size (25, 44). FcγR activation is shown as log_2_ fold change relative to negative control. Each symbol represents one sample from one patient. CMV: Cytomegalovirus reactivation under immunosuppression; HIV: HIV infection; TBC: Mycobacterium tuberculosis infection; Influenza: influenza virus infection; TX: solid organ transplantation. Solid black lines indicate the median.

**Figure 4-figure supplement 2.**
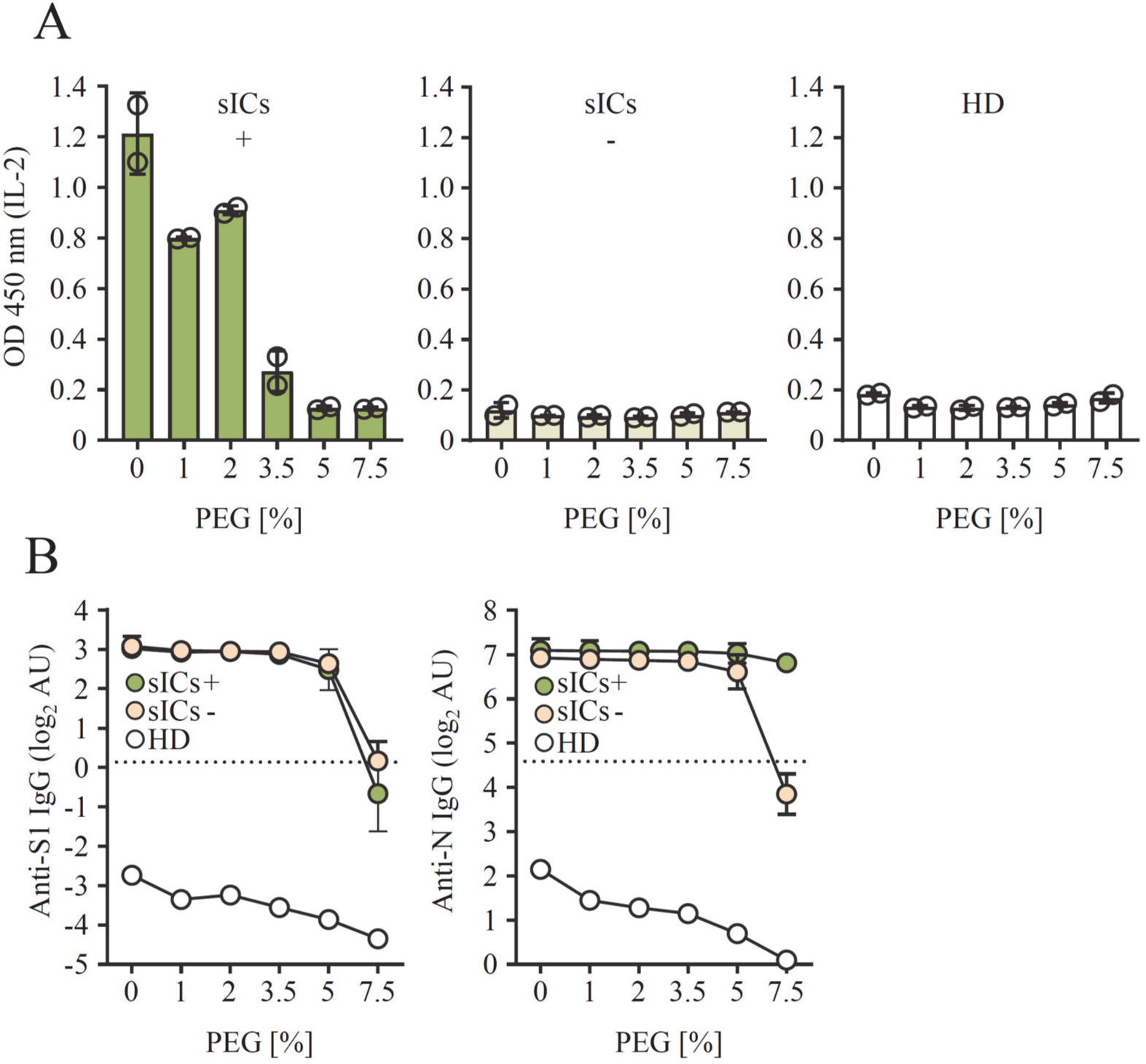
PEG precipitation eliminates sIC-mediated CD16 activation. Pools of 8 sera were incubated with equal volumes of PEG8000 to reach the indicated final PEG concentrations. A) CD16 activation after PEG-precipitation in the pool supernatant, showing either high (sICs+) or no (sICs-) CD16 activation. Sera from healthy donors (HD) were included as a negative control. Activation levels are expressed as IL-2 levels (OD 450 nm) released by reporter cells. The mean and SD of two independent experiments is depicted. B) Anti SARS-CoV-2 IgG levels against S1 (left panel) or N (right panel) IgG following PEG precipitation. The mean and SD of two independent experiments (sICs+/sICs-) is depicted.

**Figure 4-figure supplement 3.**
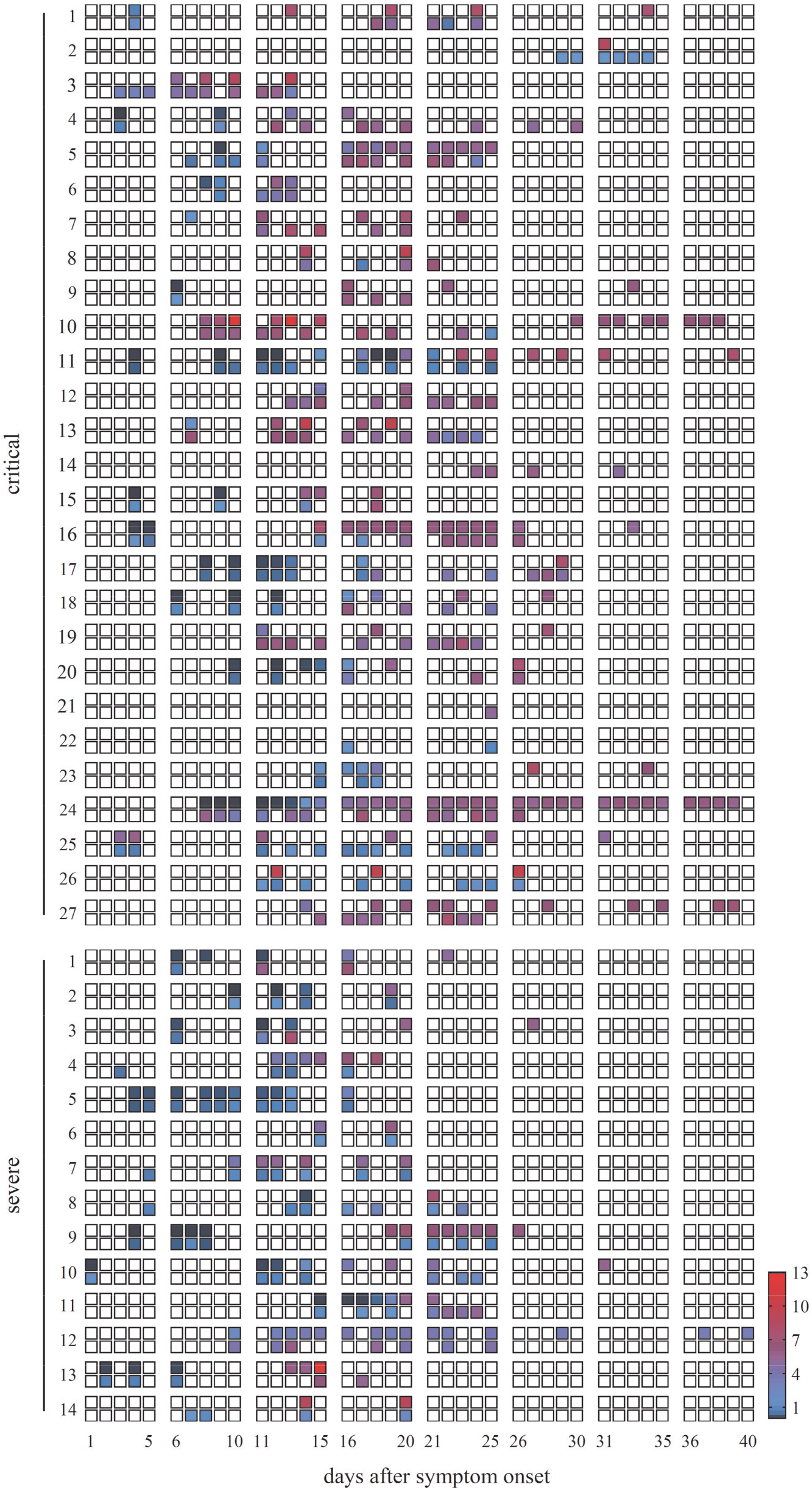
Individual CD16 activation by sICs and anti-S1 ELISA IgG kinetics post symptoms onset. Individual sera from either critically (n = 27) or severely (n = 14) diseased patients were analyzed via ELISA [AU] for anti S1-IgG (upper row) and for CD16 activation by soluble immune complexes (lower row, relative CD16 activation depicted as fold increase to the negative control) over time (1-40 days post symptom onset). White squares: not tested.

**Figure 4-figure supplement 4.**
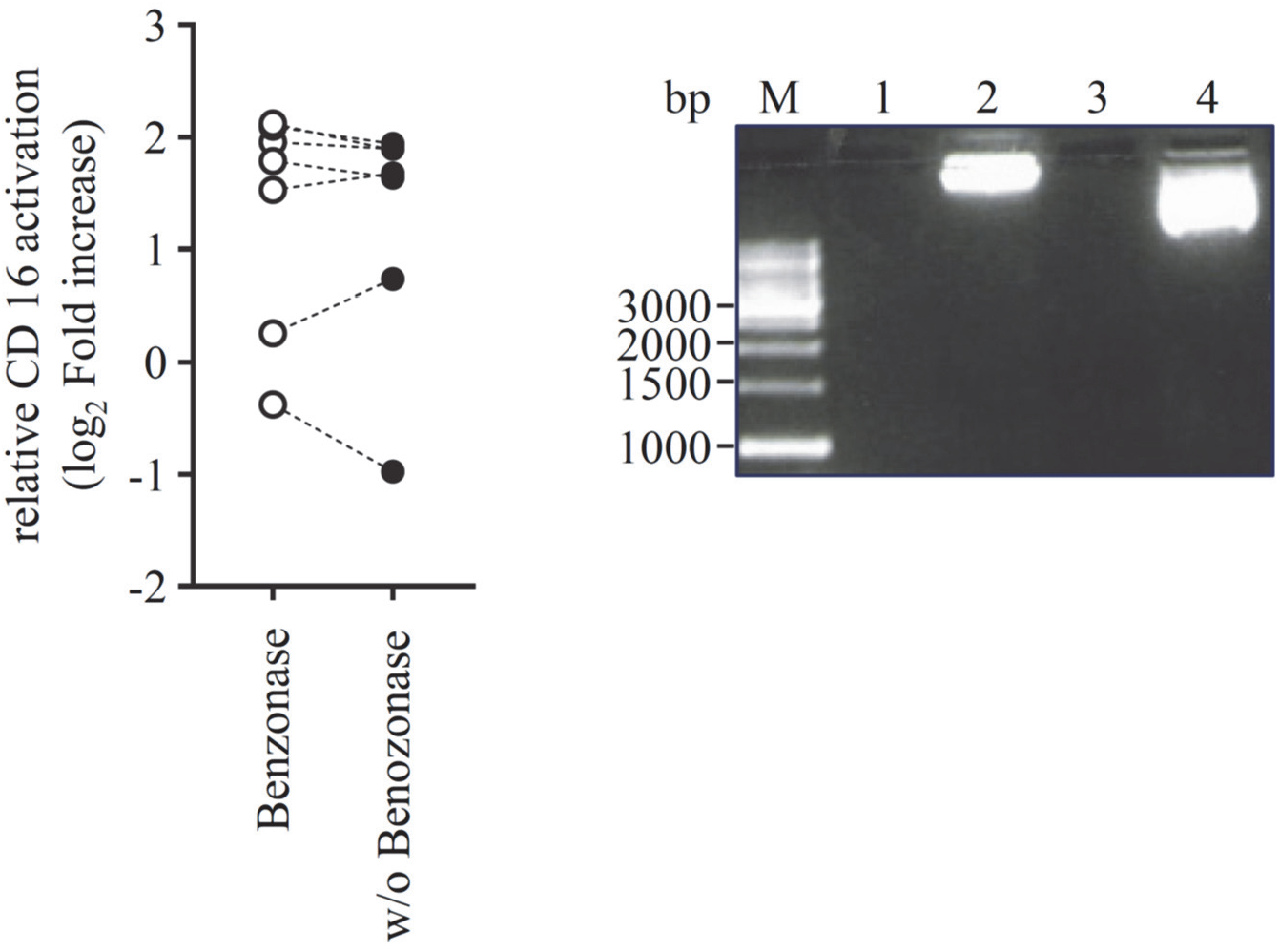
Benzonase treatment of sIC-reactive sera does not abolish CD16 activation. Left panel: sIC-mediated CD16 reactivity expressed as log_2_ fold increase to the negative control, in serum of six individual patients before and after treatment with 250 Units of Benzonase Nuclease. Right panel: As positive control, 3 μg plasmid DNA was digested. M: 1kb DNA ladder, Lane 1: benzonase digestion in the presence of human serum, lane 2: plasmid DNA w/o benzonase in the presence of human serum, lane 3: benzonase digestion in medium only and lane 4: plasmid DNA w/o benzonase in medium only.

**Figure 4-figure supplement 5.**
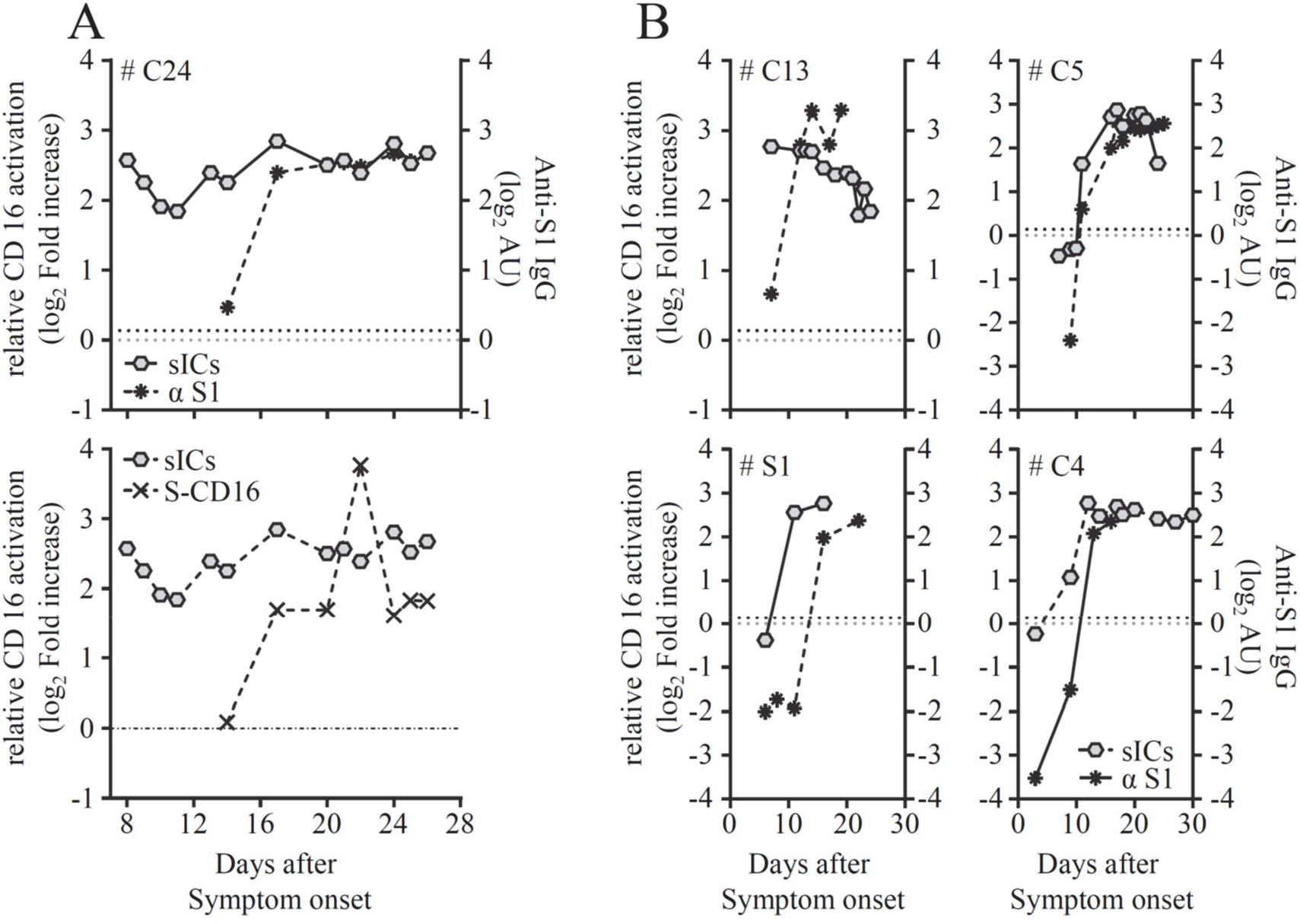
sIC formation precedes SARS-CoV-2-IgG response. Individual patients for which enough material was available were analyzed over time. A) CD16 activation by sICs-vs. S1-ELISA (top panel) and sIC –CD16 activation vs. anti-S-CD16 activation (bottom panel) in one critically ill patient, #C24. B) sIC-CD16 activation vs. S1-ELISA in four individual patients #C13, #C5, #S1 and #C4. Dashed line (black) represents commercial S1-ELISA-Cut-off level, whereas dashed line (grey) is set to 0. Individual longitudinal courses correspond to patients depicted in figure 4 supplement 3.

